# Semaphorin 6A in Retinal Ganglion Cells Regulates Functional Specialization of the Inner Retina

**DOI:** 10.1101/2023.11.18.567662

**Authors:** Rebecca E. James, Natalie R. Hamilton, Lola Nicole Huffman, Jeroen Pasterkamp, Loyal A. Goff, Alex L. Kolodkin

## Abstract

To form functional circuits, neurons must settle in their appropriate cellular locations and then project and elaborate neurites to contact their target synaptic neuropils. Laminar organization within the vertebrate retinal inner plexiform layer (IPL) facilitates pre- and postsynaptic neurite targeting, yet, the precise mechanisms underlying establishment of functional IPL subdomains are not well understood. Here we explore mechanisms defining the compartmentalization of OFF and ON neurites generally, and OFF and ON direction-selective neurites specifically, within the developing IPL. We show that semaphorin 6A (Sema6A), a repulsive axon guidance cue, is required for delineation of OFF versus ON circuits within the IPL: in the *Sema6a* null IPL, the boundary between OFF and ON domains is blurred. Furthermore, Sema6A expressed by retinal ganglion cells (RGCs) directs laminar segregation of OFF and ON starburst amacrine cell (SAC) dendritic scaffolds, which themselves serve as a substrate upon which other retinal neurites elaborate. These results demonstrate for the first time that RGCs, the first neuron-type born within the retina, play an active role in functional specialization of the IPL.

Retinal ganglion cell-dependent regulation of OFF and ON starburst amacrine cell dendritic scaffold segregation prevents blurring of OFF versus ON functional domains in the murine inner plexiform layer.

## INTRODUCTION

Elaboration of complex neural circuitry is facilitated by restricted synapse formation between distinct neuronal cell types within specific layers of laminated regions in the nervous system. During vertebrate development, lamination of early born neurons can provide a scaffold upon which later born neurons elaborate their neurites. The vertebrate retina includes a limited number of cell types, including bipolar cells (BPs), amacrine cells (ACs) and retinal ganglion cells (RGCs) which establish select synaptic connections in the inner plexiform layer (IPL) that are distributed among ∼10 sublaminae. Understanding of the development of retinal laminar organization has been greatly aided by work on direction-selective (DS) circuits, which respond to image motion through the activation of direction-selective ganglion cells (DSGCs) tuned to stimuli that move in particular directions.

DSGC dendrites stratify within the IPL, which is divided into OFF and ON subregions reflecting responses to the diminution or enhancement of illumination, respectively (Zhang et al., 2017). DSGCs tuned to fast motion in the four cardinal directions that have dendrites stratifying in both OFF and ON subregions, the sublayers S2 (OFF) and S4 (ON), respectively, and are called On-Off DSGCs (ooDSGCs; Dhande et al., 2015). DSGCs tuned to slower motion that stabilize images on the retina, termed On DSGCs (oDSGCs), are part of the accessory optic system (AOS) and have dendrites stratifying only in the ON S4 sublayer of the IPL (Hamilton et al., 2021). Projections from select classes of BPs and ACs connect with DSGCs in S2 and S4 of the IPL, comprising the DS circuits. Asymmetrically localized inhibitory synapses formed between starburst amacrine cells (SACs) and DSGC dendrites are critical for DS responses; there are both OFF- and ON-SACs, and these subtypes stratify their dendrites in either S2 or S4, respectively. In the mouse, SACs are among the first neurons to elaborate neurites within the IPL (Stacy and Wong, 2003), and as their dendritic arbors refine they form the S2 and S4 IPL sublaminae. These sublaminae serve as a scaffold that instructs subsequent DSGC and BP neurite stratification (Ray et al., 2018; Duan et al, 2018; Peng et al., 2017; Stacy and Wong, 2003).

Several molecular cues contribute to the development of murine DS circuits. SAC cell body settling in the inner nuclear layer (INL; OFF-SACs) or the ganglion cell layer (GCL; ON-SACs) requires the cell-surface protein MEGF10, as does subsequent formation of mosaic SAC cell body spacing in both the INL and GCL (Kay et al., 2012). In SACs, homotypic dendritic process self-avoidance requires protocadherins (Lefebvre et al., 2012), and classical cadherins and contactin cell adhesion molecules are important for select BP and DSGC targeting of axons and dendrites, respectively, to S2 and S4 (Duan et al., 2014; Duan et al., 2018; Peng et al., 2017). Further, interactions between the fibronectin leucine-rich repeat transmembrane 2 (FLRT2) receptor and transmembrane UNC5 ligand abrograte FLRT2-latrophilin (LPHN) adhesive interactions, thereby regulating DSGC dendritic targeting to correct INL strata (Prigge et al., 2023). In addition, the transmembrane protein semaphorin 6A (Sema6A) and its plexin A2 (PlexA2) receptor also play key roles in DS circuit development. Sema6A is enriched in ON regions of the IPL (Matsuoka et al, 2011a), and *Sema6A* loss-of-function (LOF) leads to mis-stratification of SAC dendrites, disruption of ON-SAC dendritic arbor morphology, and loss of ON but not OFF ooDSGC DS tuning responses (Sun et al., 2013). However, the cell types in which Sema6A is required were not previously determined owing to use thus far of a global *Sema6a* LOF allele.

Here, we revisit Sema6A/PlexA2 signaling in DS circuit and overall retinal development using newly generated mouse mutants allowing for cell type-specific assessments of Sema6A function. We refine the role played by Sema6A in SAC development, and define roles for Sema6A forward and reverse signaling in distinct aspects of DS circuit formation. Analysis of Sema6A function in RGCs reveals novel roles for RGCs in directing SAC dendrite lamination and delimiting the ON regions of the IPL. These results provide insight into the development of retinal laminar organization.

## RESULTS

### *Sema6a* is not Required in SACs for Correct Dendritic Arbor Lamination

We generated an HA-epitope knock-in *Sema6a* conditional allele (*Sema6a^HA-F^*) that produces an N-terminal, HA-tagged, fully functional Sema6A protein (HA-Sema6A) in the absence of Cre recombinase to investigate the cell-type-specific requirements for *Sema6a* in SAC development (Fig. 1A; Fig. S1A-D). HA-Sema6A expression mirrors that of endogenous Sema6A, assessed by comparing HA immunoreactivity to an antibody directed against Sema6A (Kerjan et al., *2*005) in postnatal day 10 (P10) retinal sections (Fig. 1B). This Sema6A antibody can be unreliable, often yielding high background with a speckled, non-specific signal that fails to recapitulate wildtype Sema6A staining *in vivo* and *in vitro* (Fig. S2), complicating a thorough assessment of Sema6A expression. However, HA-Sema6A immunolabeling faithfully and reproducibly reports Sema6A expression (Figures 1B), providing a robust avenue for analysis of Sema6A protein distribution.

**Fig. 1.**
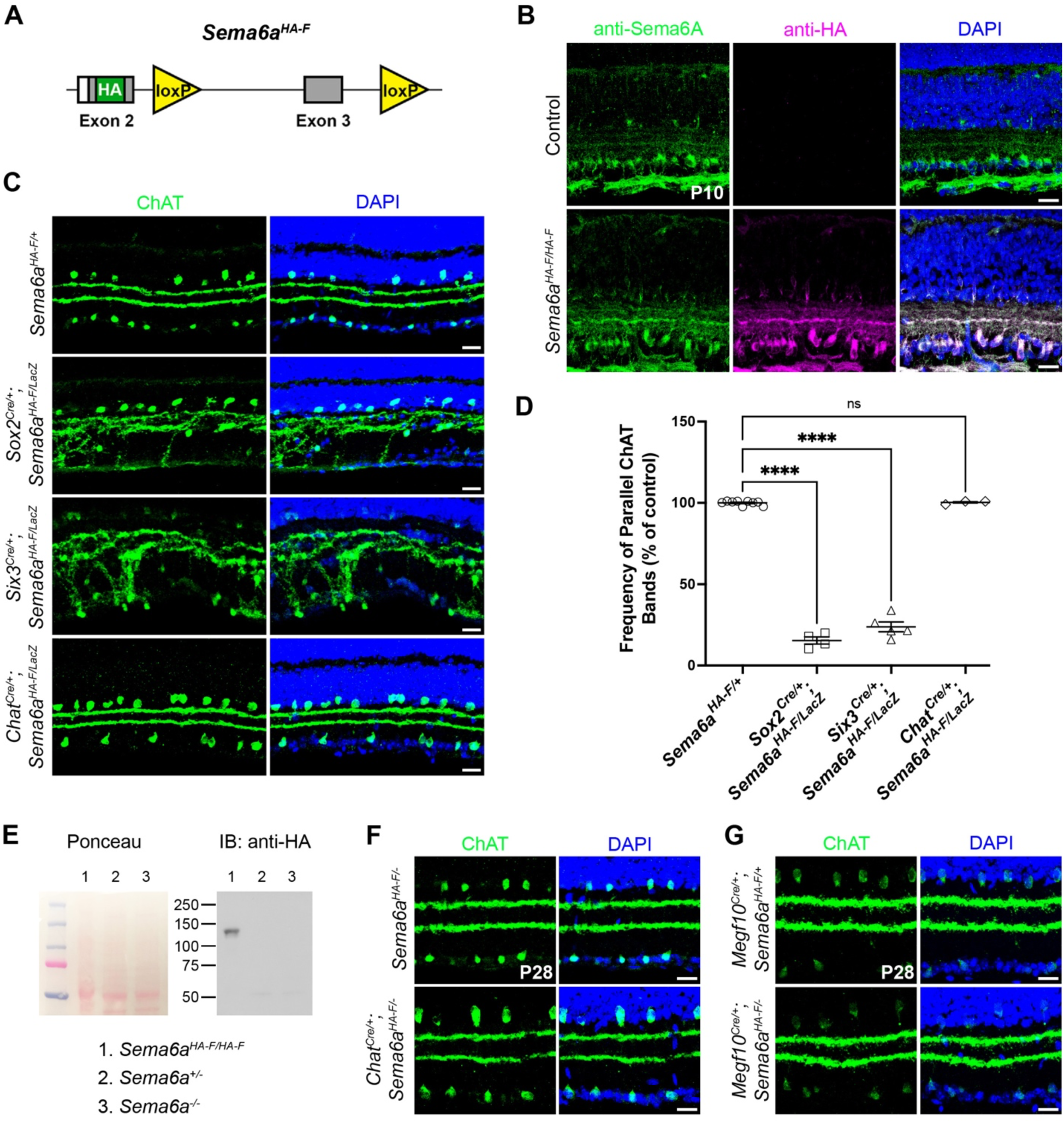
SAC-derived Sema6A does not contribute to layer-specific targeting of SAC dendrites. **(A)**Schematic of the *Sema6a* conditional allele design. Cre-mediated loxP recombination generates a frameshift and premature stop codons. **(B)** Anti-Sema6A (green) in P10 control and *Sema6a^HA-F/HA-F^*retinas. Anti-HA (magenta) is specific and recapitulates anti-Sema6A (green). **(C)** SAC (anti-ChAT) dendrite lamination for indicated genotypes. **(D)** Quantification of the frequency of segregated, parallel ChAT^+^ SAC strata in the IPL. **(E)** Western blot showing HA-Sema6A (∼130 kDa) in *Sema6a^HA-F/HA-F^* but not *Sema6a^+/-^* or *Sema6a^-/-^* (null allele derived from germline-recombined *Sema6a^HA-F^*) in cortical lysates. Ponceau S, loading control. **(F)** Neonatal (*Chat^Cre^*) SAC-specific *Sema6a* cKO. **(G)** Embryonic (*Megf10^Cre^*) SAC-specific *Sema6a* cKO. Scale bars, 20μm.

We first crossed our new *Sema6a^HA-F^*allele to a germline Cre driver (*Sox2^Cre^*; Hayashi et al., 2002) in a genetic background heterozygous for *Sema6a^HA-F^* and the *Sema6a* gene trap LOF allele (*Sema6a^LacZ^*) (Leighton et al., 2001, Matsuoka et al., 2011a, Sun et al., 2013). Unexpectedly, germline *Sema6a* LOF resulted in more severe SAC lamination defects than those previously observed (Sun et al., 2013; Fig. S1A), consisting of a nearly complete fusion of the SAC IPL dendritic stratifications (Fig. 1C) that is more pronounced than the crossovers observed in *Sema6a^LacZ^* mutants (Sun et al., 2013). Therefore, the *Sema6a^LacZ^* allele is hypomorphic. This phenotype was also evident in the central retina of *Six3^Cre/+^* (Furuta et al., 2000) conditional knockout (cKO) retinas, where Cre is strongly expressed in central but not peripheral retinal progenitors (Ray et al., 2018). However, when *Sema6a* was selectively removed from postmitotic SACs using *Chat^Cre^*(Rossi et al., 2011), SAC dendrites were appropriately segregated and SAC cell bodies were normally distributed (Fig. 1C,D). HA- Sema6A is not produced in *Sema6a^+/-^* or *Sema6a^-/-^*animals harboring our newly generated *Sema6a* null allele (Fig. 1E).

We examined SAC segregation in *Chat^Cre^; Sema6a^HA-F^*^/-^ mice, which are heterozygous for the new *Sema6a* null allele instead of *Sema6a^LacZ^*, and found that SAC IPL lamination was similarly unaffected (Fig. 1F). Since expression of *Chat^Cre^* coincides with SAC dendrite lamination (Ray et al., 2018), we asked if use of *Megf10^Cre^*, which is expressed exclusively in SACs embryonically (as early as E16.5; Peng et al., 2020), revealed a SAC lamination defect. Embryonic removal of *Sema6a^HA-F^*using *Megf10^Cre^* also did not impact SAC lamination (Fig. 1G). These results demonstrate that Sema6A expressed by SACs is not required for OFF/ON-SAC dendrite segregation.

### *Sema6a* in SACs Promotes Dendrite Outgrowth and Self-avoidance

Sema6A and PlexA2 contribute to ON-SAC dendritic arbor radial symmetry (Sun et al., 2013), suggesting that isoneuronal *trans* repulsive signaling promotes sister dendrite self-avoidance (Fig. S3A). Therefore, we examined SAC morphology in *Sema6a* null retinas at P14 by recombining the fluorescent reporter *ROSA^LSL-TdTomato^* (Madisen et al., 2010) in sparse SACs using *ChatCre^ER^* (Rotolo et al., 2008). In *Sema6a^LacZ^* mutants, OFF-SACs developed appropriately, while *Plxna2^-/-^* OFF-SACs exhibited a modest defect in dendritic outgrowth (Sun et al, 2013). However, *Sema6a^-/-^*OFF-SAC outgrowth was reduced by 45% and OFF-SAC symmetry was also modestly diminished (Fig. 2A,B). ON-SAC outgrowth and symmetry were more severely disrupted in *Sema6a^-/-^* compared to *Sema6a^LacZ^* retinas (Fig. 2A,C; Sun et al., 2013). For both OFF- and ON-SACs, these outgrowth defects were accompanied by robust distal dendrite self-avoidance errors (Fig. 2A).

**Fig. 2.**
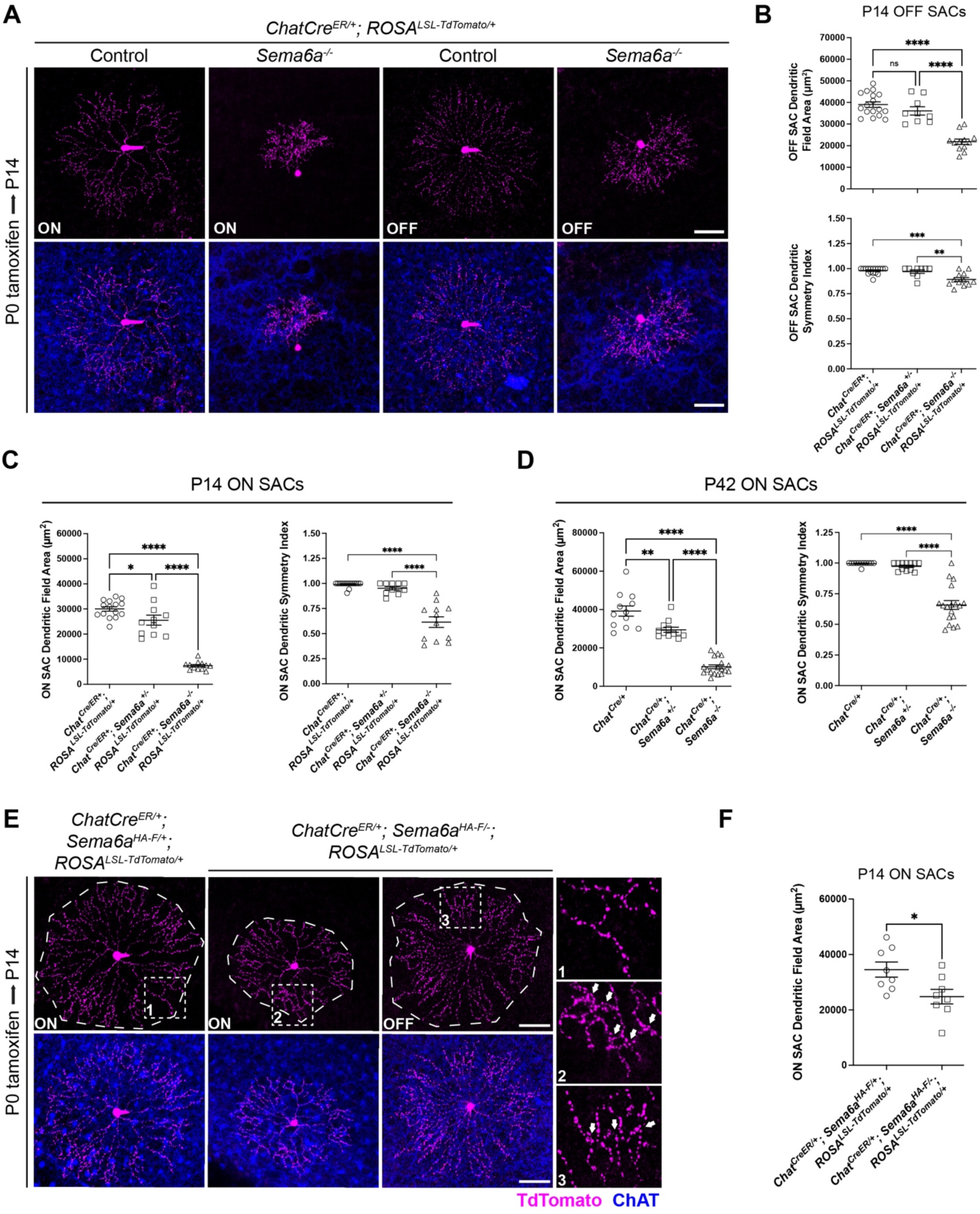
Sema6A in SACs promotes neurite outgrowth and distal dendrite self-avoidance. **(A)**P14 SACs labeled by recombination of *LSL-TdTomato* by P0 tamoxifen injection. Control is *Sema6a^+/+^*. Quantification of P14 OFF **(B)** and ON **(C)**, and P42 ON **(D)** SAC dendritic arbor area and symmetry. P42 ON-SACs quantified in **(D)** labeled by AAV2-Flex-GFP intraocular injection at P28. **(E)** P14 ON- and OFF-SACs labeled by P0 tamoxifen administration exhibit distal dendrite self-avoidance errors. **(F)** Quantification of dendritic field area following cell-autonomous removal of *Sema6a* in isolated P14 ON-SACs. Scale bars, 50μm.

We next examined isolated SAC morphology using a Cre-dependent *AAV2-FLEX-GFP* virus. GFP^+^ ON-SAC dendrite development was severely impaired in *Chat^Cre/+^; Sema6a*^-/-^ mutants, demonstrating reduced dendritic outgrowth, diminished dendritic process self-avoidance, and disrupted dendritic arbor radial symmetry (Fig. S3B). The remaining branches of *Sema6a^-/-^* null ON-SAC arbors were often aberrantly bundled (Fig. S3B, arrow). *Sema6a^+/-^*ON-SAC dendrite outgrowth (Fig. 2D) and distal dendrite self-avoidance (Fig. S3B, arrowheads) are impaired, reminiscent of *Sema6a^LacZ/LacZ^*mutant ON-SAC arbors, demonstrating that *Sema6a* is haploinsufficient for ON-SAC dendritic arbor development. The AAV reporter cannot penetrate deeply enough to sufficiently label sparse OFF-SACs, precluding quantification of OFF-SAC phenotypes. Further, SAC strata are effectively fused in *Sema6a^-/-^* retinas, so we were unable to examine general ON- versus OFF-SAC dendritic plexus organization (see Fig. 1 and Fig. 5).

We next performed sparse *Sema6a* cKO using *ChatCre^ER^* and *ROSA^LSL-TdTomat^*° to assess whether cell-autonomous Sema6A signaling promotes self-avoidance of branching sister dendrites (Fig. S3A; see also: Sun et al., 2013). HA-Sema6A and PlexA2 are both enriched in the distal one-half of cultured SAC dendrites, accumulating at branch points (Fig. S4). Targeted sparse *Sema6a* cKO ON-SACs exhibited distal dendrite crossovers and impaired dendritic arbor outgrowth (Fig. 2E; Sun et al., 2013), demonstrating that *Sema6a* cell-autonomously promotes self-avoidance and outgrowth of ON-SAC dendrites. Disrupted distal dendrite self-avoidance was also observed in all targeted sparse OFF-SACs (Fig. 2E), showing that isoneuronal *trans* repulsive signaling between Sema6A and PlexA2 promotes dendrite self-avoidance in all SACs. Unexpectedly, cell-autonomous removal of *Sema6a* in individual SACs did not alter SAC symmetry (Fig. 2E), and cell-type conditional removal of *Sema6a* in all SACs did not disrupt SAC radial symmetry or plexus organization (Fig. S3C). Instead, only distal dendrite self-avoidance was disrupted by conditional cell-autonomous removal of *Sema6a* from SACs.

Together, these data demonstrate that large gaps in ON-SAC plexus organization observed in *Sema6a* mutant retinas (Fig. S3B,D, arrowheads; Sun et al., 2013) do not result from the distal dendrite self-avoidance defects that arise when SACs no longer express Sema6A. Further, Sema6A acts in all SACs to promote dendrite outgrowth and self-avoidance, suggesting that perturbations in ON-SAC symmetry and plexus organization in *Sema6a* mutants arise through a SAC non-autonomous mechanism.

### Sema6A is Expressed by ON and OFF SACs

Upon close examination of HA-Sema6A expression, HA-Sema6A was not obvious in ON-SACs at P4 and P10 (Fig. 3A, arrowheads), contrary to our expectations. However, robust HA- Sema6A signal from RGCs could preclude detection of Sema6A in SACs in retinal cross sections, so we examined HA-Sema6A in P10 whole mount retinas. ON-SACs imaged through the center of the GCL did not appear to express HA-Sema6A; however, many RGCs labeled by the pan-RGC marker RBPMS (Rodriguez et al., 2014) expressed HA-Sema6A (Fig. 3B, top). Biasing our imaging plane toward the IPL revealed that the very same ON-SACs (Fig. 3B, top: white dashed circles) that appeared devoid of HA-Sema6A were indeed HA-Sema6A^+^ (Fig. 3B, middle). Using this same imaging paradigm for OFF-SACs we similarly detected HA-Sema6A (Fig. 3B, bottom), in line with *Sema6a* cKO OFF-SACs exhibiting outgrowth and distal dendrite self-avoidance defects (Fig. 2D).

**Fig. 3.**
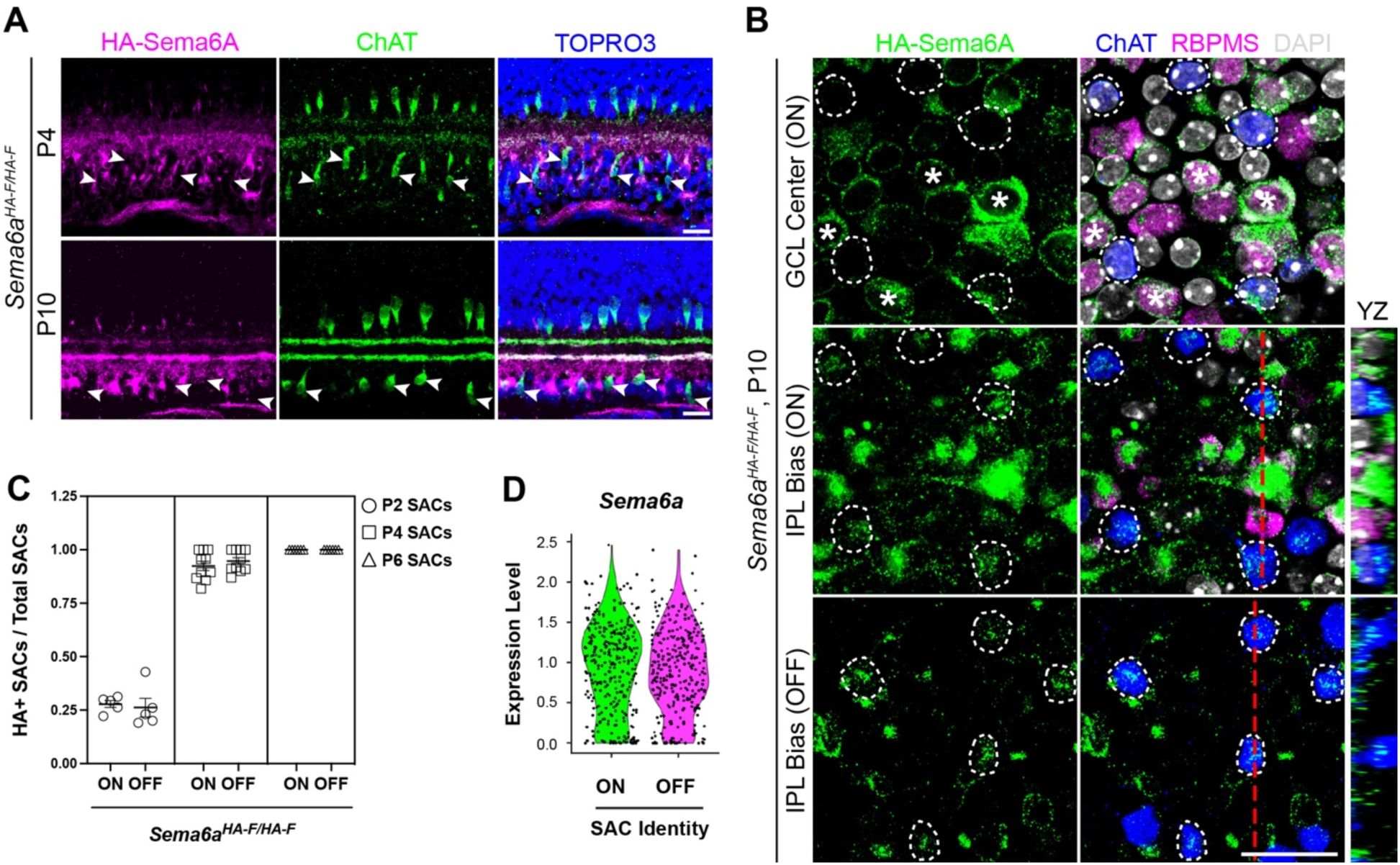
Sema6A is expressed by both ON- and OFF-SACs. **(A)**HA-Sema6A (magenta) is not evident in P4 and P10 *Sema6a^HA-F/HA-F^*ON-SACs (arrowheads). **(B)** HA-Sema6A (green) is not evident in ChAT^+^ ON-SACs (dashed circles) when imaging through the GCL center (top) in whole mount retinas, but is in the same SACs (dashed circles) when the imaging plane is shifted toward the IPL in the z-axis (middle). HA-Sema6A is detected in RBPMS^+^ RGCs (asterisks). HA-Sema6A is also detected in OFF-SACs (bottom, white circles) using this IPL- biased imaging strategy (YZ orthogonal views: red dashed lines, right). **(C)** Quantification HA- Sema6A^+^ SACs across early postnatal development. **(D)** Violin plot of *Sema6a* expression in SACs isolated at P1, P4, P8, and P11. Scale bars, 20μm.

We next characterized Sema6A expression across early postnatal development and found that only ∼25% of ON- and OFF-SACs expressed Sema6A protein at P2, when SAC lamination is complete (Ray et al., 2018; Sun et al., 2013). Between P2 and P4, Sema6A protein expression in SACs inceases, and by P6 all SACs express Sema6A (Fig. 3C). We previously observed an apparent lack of Sema6A expression in some SACs cultured *in vitro* and in OFF-SACs in retinal cross sections (Sun et al. 2013), yet here we detect HA-Sema6A protein in all SACs. We attribute this to the unreliability of the antibody directed against Sema6A used in previous studies (Sun et al., 2013), leading us to favor the use of *Sema6a^HA-F^* to report Sema6A expression.

To resolve the timing and cell types that express *Sema6a* with temporal resolution, we generated a new *Sema6a^CreER^*knock-in allele (Fig. S5A; Materials and Methods). *Sema6a^CreER^*- driven expression of *ROSA^LSL-TdTomato^* was evident in ∼20% of both all SACs from P1-P3, while ∼70% of RGCs were TdTomato^+^ (Fig. S5B,C). *Sema6a* was also expressed in ∼36% of presumptive displaced non-SAC amacrine cells (arrows, Fig. S5B). *Sema6a* transcriptional activity in SACs assessed with *Sema6a^CreER^* peaked at ∼40% by P8 and was nearly absent by P14 (Fig. S5D).

We confirmed these expression results using an adapted Smart-seq2 (Chevee et al, 2018) single-cell mRNA sequencing (scRNA-seq) protocol that facilitates deep sequencing to discern gene expression differences between closely related OFF- and ON-SACs. FACS-isolated TdTomato^+^ SACs from P1, P4, P8, P11, and P16 *Chat^Cre/+^; ROSA^LSL-TdTomato/+^* retinas were profiled (see Methods; >425,000 reads/cell, >3900 genes/cell for P1-P11 libraries). SACs at each postnatal timepoint, except P16, unbiasedly clustered into two groups, identified by *Fezf1* and *Rnd3* as ON- or OFF-SACs respectively (Peng et al., 2020; Fig. S6A). All clusters were enriched for genes with known SAC expression (Fig. S6B-E). *Sema6a* expression was distributed across all SACs and SAC *Sema6a* and *Plxna2* expression levels were comparable across postnatal development and were significantly downregulated by P16 (Fig. 3D, Fig. S6F- I); this is consistent with *Sema6a^CreER^* reporter activity turning off by P14 (Fig. S5D). By P16, genes contributing to the formation and specialization of ON- versus OFF-SAC circuits (*Fezf1* and *Rnd3*) were downregulated (Fig. S6A,E) whereas genes devoted to synaptic function (*Slc18a3*, *Gad1*, and *Synpr*) were upregulated (Fig. S6C,D,J), suggesting that once specialized ON- and OFF-SAC circuits have formed, the differentially-expressed genes underlying that specialization turn off and SAC subtypes become more similar to one another.

Together, these results support a cell-autonomous function for Sema6A in ON- and OFF-SAC dendritic arbor development, but not lamination. Furthermore, since (1) SAC dendritic stratification is complete by P1 (Ray et al., 2018), (2) *Sema6a* is expressed by few SACs at this time (∼20%), and (3) Sema6A is robustly expressed by other cell types in the neonatal retina (Fig. 3, Fig. S5), another cell type must promote Sema6A-mediated SAC dendritic stratification.

### Sema6A Forward Signaling Through PlexA2 Directs SAC Dendritic Arbor Elaboration

Sema6A can function as a ligand (forward signaling) via PlexA2 or as a receptor (reverse signaling; Battistini & Tamagnone, 2016). To determine whether reverse Sema6A signaling influences SAC development, we used a *Sema6a^ΔCyt^* allele designed to conditionally remove the Sema6A cytoplasmic domain upon Cre expression (Verhagen et al., 2023). We first examined organization of the ON-SAC dendritic plexus as a proxy for ON-SAC radial symmetry, since the ON-SAC plexus has gaps only when SAC radial symmetry is perturbed (note ON-SAC plexus uniformity, Fig. S3C, where radial symmetry is preserved, versus Fig. S3D; see also: Sun et al., 2013). Conditional removal of the Sema6A cytoplasmic domain in SACs did not alter SAC plexus uniformity: large gaps indicative of impaired SAC radial dendritic symmetry were not observed in the *Megf10^Cre/+^; Sema6a^ΔCyt^* SAC plexuses (Fig. S7A,B). Lamination of the SAC strata in *Sema6a^ΔCyt^* cKO retinas was also preserved (Fig. S7C). Together, these results show that reverse Sema6A signaling in SACs is dispensable for SAC dendritic arbor development.

PlexA2, a receptor for forward repulsive Sema6A signaling (Pasterkamp, 2007), is exclusively expressed in SACs in the neonatal retina and is required for SAC dendrite lamination and development (Matsuoka et al, 2011; Sun et al. 2013). Plexin forward signaling requires plexin cytoplasmic domain RasGAP activity (Pascoe et al., 2015). We used the *Plxna2^R1746A^* allele (Zhao et al., 2018), in which the PlexA2 RasGAP activity is abolished by a point mutation, (called here *Plxna2^ΔRasGAP^*) to ask if PlexA2 forward repulsive signaling impacts SAC development. *Plxna2^+/-^* retinas rarely exhibit SAC strata crossovers (0.39 ± 0.17 per mm). There were significantly more crossovers (4.17 ± 0.67 crossovers per mm) in *Plxna2^ΔRasGAP/-^* retinas, but this was only ∼27% of *Plxna2^-/-^* retinas (15.63 ± 0.97 crossovers per mm), suggesting that PlexA2 RasGAP domain-mediated forward signaling only partially regulates SAC lamination (Fig. 4A, B).

**Fig. 4.**
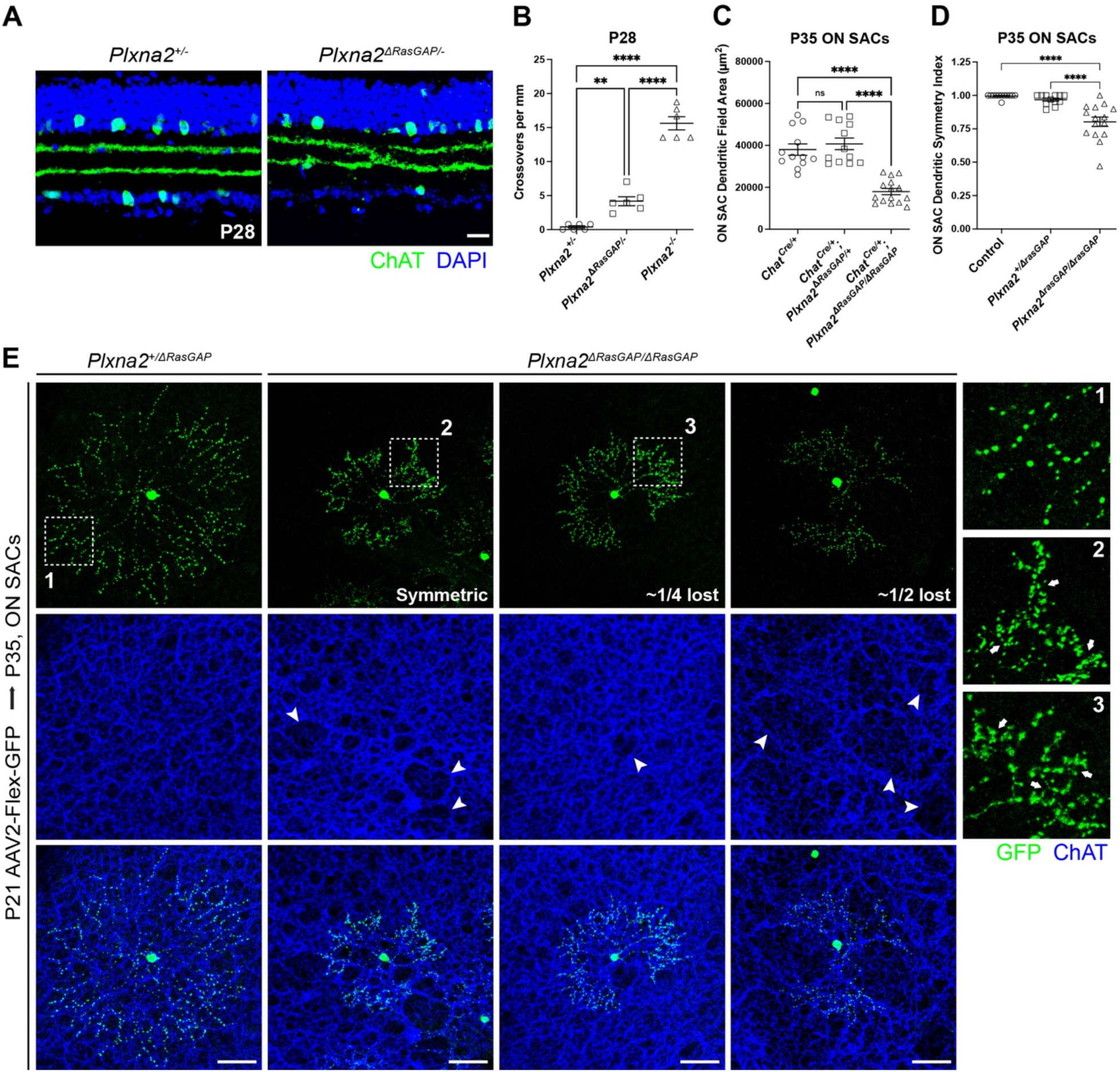
Forward signaling through the PlexA2 RasGAP domain is required for SAC development. **(A)**SAC lamination (anti-ChAT) in P28 *Plxna2^τ<RasGAP/-^* retinas. Scale bar, 20µm. **(B)** Quantification of crossovers in *Plxna2^τ<RasGAP/-^* vs. *Plxna2^-/-^*mutants. Quantification of P35 AAV2-Flex-GFP labeled ON-SAC dendritic field area **(C)** and symmetry **(D)**. **(E)** Representative P35 AAV2-Flex-GFP labeled ON-SACs. Varying degrees of ON-SAC radial symmetry are observed in *Chat^Cre/+^; Plxna2^11RasGAP/11RasGAP^* retinas. Distal dendrites of every *Plxna2^11RasGAP^* mutant ON-SAC examined exhibited self-avoidance errors. The *Plxna2^11RasGAP^* mutant ON-SAC plexus contains large holes and gaps (arrowheads). Scale bar, 50µm.

We next examined SAC morphology in *Chat^Cre/+^; Plxna2^ΔRasGAP^* mutant retinas transduced with AAV-Flex-GFP. *Plxna2^ΔRasGAP/ΔRasGAP^* (and *Plxna2^ΔRasGAP/-^*; *data not shown*) mutant ON-SAC dendrite outgrowth was severely stunted; ON-SAC arbors displayed symmetry and distal dendrite self-avoidance defects, comparable to *Sema6a^-/-^* and *Plxna2^-/-^* mutant ON-SACs (Fig. 4C-E; Sun et al., 2013). Examination of the ON-SAC dendritic plexus in *Plxna2^ΔRasGAP/ΔRasGAP^*mutants also revealed large gaps, further demonstrating the link between ON-SAC radial symmetry and plexus organization (Fig. 4E, arrowheads). These results show that forward signaling through PlexA2 is required for ON-SAC dendrite outgrowth, arbor symmetry, and self-avoidance. Few OFF-SACs were sufficiently labeled to permit quantification of OFF-SAC dendritic arbor morphology. Those that were imageable exhibited radial dendritic arbors with distal dendrite self-crossovers (Fig. S7D), strongly suggesting that forward signaling through PlexA2 is required for self-avoidance in all SACs. Large gaps in the OFF-SAC dendritic plexus were not detected in *Plxna2^ΔRasGAP/ΔRasGAP^*mutants (Fig. S7E), distinguishing between Sema6A/PlexA2-dependent SAC distal dendrite self-avoidance versus overall dendritic arbor symmetry.

### Sema6A is Broadly Required for Inner Retina Development

SAC dendrites are scaffolds that direct lamination of incoming retinal neurites (Stacy and Wong, 2003; Peng et al., 2017; Duan et al., 2018). Our present results demonstrate that previous studies (Matsuoka et al., 2011a; Sun et al., 2013) were conducted in a hypomorphic background. Therefore, we reexamined inner retinal phenotypes in *bona fide Sema6a^-/-^* retinas. As expected (Fig. 1 and Fig. 2), the SAC strata defined by vesicular acetylcholine transporter (VAChT) immunolabeling were fused in *Sema6a^-/-^* retinas (Fig. 5A). VGlut3 expression, which labels a population of ACs (Fremeau et al., 2002) stratifying neatly between the S2 and S4 SAC layers, was no longer confined between S2-S4 in *Sema6a^-/-^* retinas (Fig. 5B). Consistent with our previous findings in the *Sema6a^LacZ^* hypomorphic background (Matsuoka et al., 2011a), tyrosine hydroxylase (TH)-expressing ACs misproject a portion of their dendritic arbor into ON IPL layers (Fig. 5A).

**Fig. 5.**
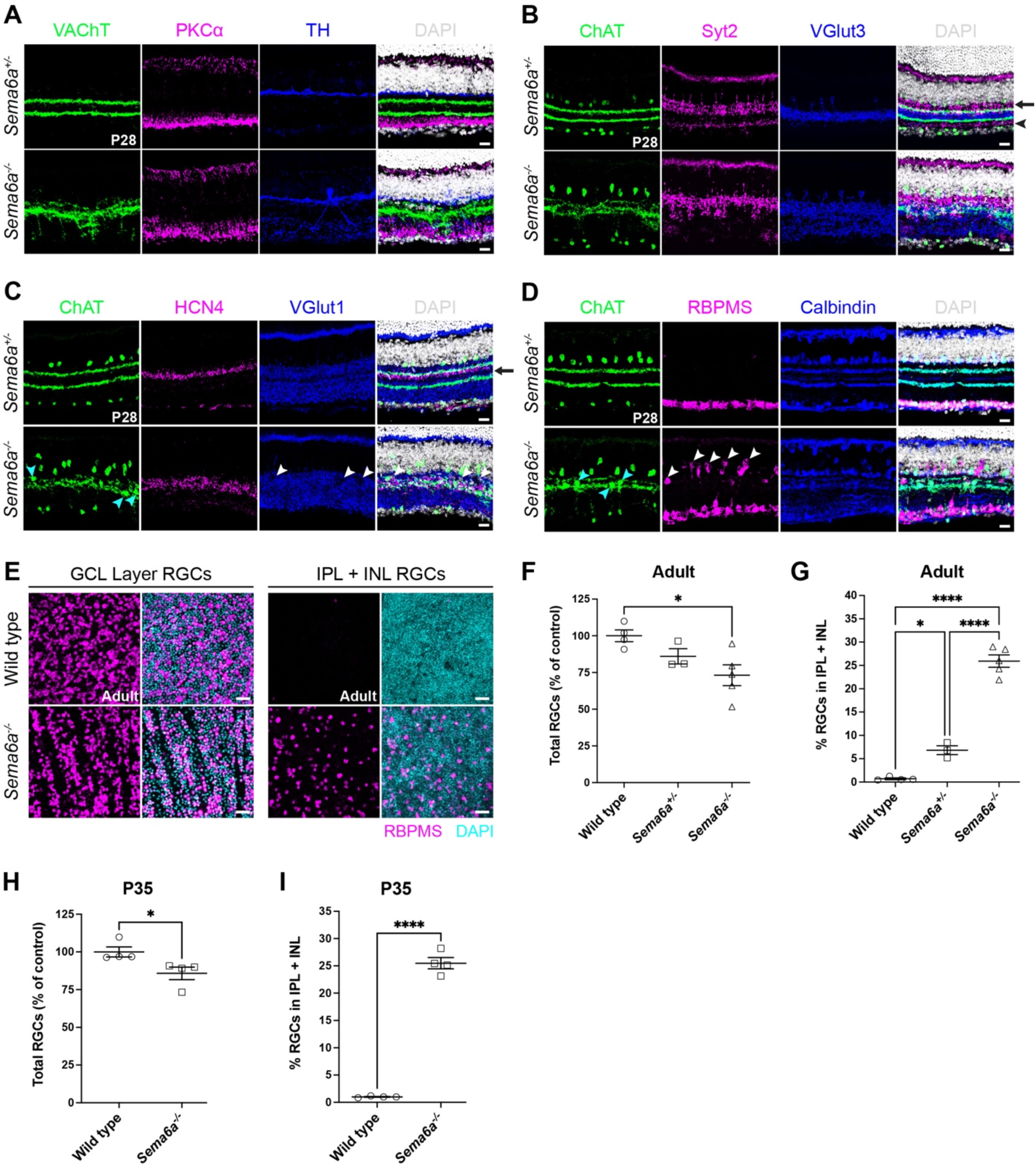
*Sema6a* is broadly required for inner retina development. P28 retina cross sections labeled with: anti-VAChT (SACs), anti-PKCɑ (rod BCs), and anti-tyrosine hydroxylase (TH^+^ ACs) **(A)**; anti-ChAT (SACs), anti-Syt2 (type 2 OFF CBCs, intense labeling indicated by arrow; type 6 ON CBCs, weak labeling indicated by arrowhead), and anti-VGlut3 (VGlut3^+^ ACs) **(B)**; anti-ChAT (SACs), anti-HCN4 (type 3a BCs), and anti-VGlut1 (BC axon terminals in the inner retina) **(C)**; and anti-ChAT (SACs), anti-RBPMS (RGCs), and anti-calbindin (subtypes of RGCs and ACs, including SACs) **(D)**. **(E)** Z-projections through the indicated regions of whole mount retinas. **(F)** Normalized quantification of total RGCs. **(G)** Quantification of IPL+INL displaced RGCs. **(H)** Normalized quantification of total RGCs at P35. *Sema6a^-/-^* mutant retinas are missing ∼15% of total RGCs. **(I)** Quantification of IPL + INL displaced RGCs at P35. Wild type, C57BL/6. Scale bars (A-D), 20µm. Scale bar (E), 40µm.

In addition, we noted a modest expansion of the PKCα rod bipolar cell (RBC) axon termination zone (Haverkamp et al., 2003), which is normally apically-delimited by the ON-SAC S4 IPL layer and basally-delimited by the GCL (Fig. 5A). We also examined projections of OFF type 2 cone bipolar cells (CBCs), which are strongly labeled by anti-synaptotagmin-2 (Syt2) in S1 and S2 (Fig. 5B, arrow), and ON type 6 CBCs, which are weakly labeled by anti-Syt2 in S5 (Fig. 5B arrowhead; Fox and Sanes, 2007; Wässle et al., 2009). In *Sema6a^-/-^*retinas, we observed clumpy regions of basally misprojecting type 2 CBCs into ON IPL layers (Fig. 5B). The type 6 ON CBC termination band was completely disrupted in *Sema6a^-/-^* retinas; instead of a distinct band of Syt2^+^ axon terminals in S5, diffuse, weak Syt2^+^ patches were observed. Similar to RBC, Syt2^+^ type 2 (OFF), and type 6 (ON) CBC axon termination zone expansion, we observed a broadening of HCN4^+^ OFF type 3a CBC (Müller et al., 2003) axon terminals in *Sema6a^-/-^* retinas. Type 3a CBCs usually terminate in a tight band partially overlapping and just basal to the S2 OFF-SAC layer (Fig. 5C). This HCN4^+^ band lies just apical to the middle of S3, a region with sparse BC axon terminations revealed by vesicular glutamate transporter-1 (VGlut1) (Johnson et al., 2003), which likely represents a boundary between OFF and ON IPL circuits (Fig. 5C, black arrow). In *Sema6a^-/-^* retinas, type 3a OFF CBC axons terminate diffusely across S2 and S3. The BC axon terminal-free, VGlut1^-^, region in S3 is no longer evident in *Sema6a^-/-^* mutants, and frequent holes, corresponding to displaced cell bodies, disrupt the VGlut1 staining pattern (Fig. 5C, white arrowheads). Together, these results suggest a blurring of the boundary separating OFF versus ON domains of the IPL in *Sema6a^-/-^* mutants.

Finally, we examined RGC and AC neurite lamination patterns in *Sema6a^-/-^* retinas using RBPMS and calbindin^+^. Like in *Sema6a^LacZ^* retinas (Matsuoka et al. 2011a), calbindin^+^ neurites misproject basally into S4/S5 in *Sema6a^-/-^* mutants (Fig. 5D). Migration of RGCs into the GCL was disrupted in *Sema6a^-/-^* retinas (Fig. 5D, white arrowheads), consistent with delayed migration of retinal progenitors in *Sema6a^LacZ^* mutants (Belle et al., 2016). Similarly, SAC cell bodies are occasionally detected in the IPL (Fig. 5C,D, cyan arrowheads). To quantify RGC displacement, we assessed apicobasal RGC cell body positioning in a centrally-positioned region of interest along the central-to-peripheral axis in each of four quadrants in wildtype, *Sema6a^+/-^*, and *Sema6a^-/-^*retinas. RBPMS^+^ RGCs in the IPL and INL were considered “displaced”. In adult *Sema6a^-/-^* retinas we observed a 26% loss of RGCs and a 25% increase in displaced RGCs (Fig. 5E-G). We examined RGC phenotypes in juvenile *Sema6a^-/-^* retinas and found that ∼15% of total RGCs were lost at P35, indicating that RGC loss increases over time (Fig. 5H). P35 mutant retinas also exhibited a 25% increase in displaced RGCs (Fig. 5I), demonstrating that RGC mislocalization remains constant and precedes RGC loss. This loss of RGCs over time likely reflects targeting defects and/or lack of trophic support. The identities of displaced/lost RGCs remain to be determined. Melanopsin^+^ intrinsically photosensitive RGCs (ipRGCs) only account for ∼2.5% of total RGCs, with displaced ipRGCs representing ∼0.35% of the population (Valiente-Soriano et al., 2014). Thus, normally displaced ipRGCs do not significantly impact our quantification of RGC displacement.

Taken together, these results show that in the complete absence of Sema6A, BPs, ACs and RGCs exhibit a broad range of defects including neurite IPL mis-stratification, cell body settling errors (RGCs and SACs), and reduced survival (RGCs). Deficiencies in cell migration (Fig. 5C- G) and neurite lamination (Fig. 5A-D) of various retinal cell types suggest distinctions between early and late roles for Sema6A in retinal development.

### Sema6A Directs Retinal Development Through Plexin-dependent and -independent Mechanisms

PlexA1-A4 are each expressed in the developing retina, yet only PlexA2 and PlexA4 are receptors for Sema6A in the retina (Matsuoka et al., 2011a,b; Sun et al., 2013). Therefore, we analyzed retinal neuron phenotypes in *Plxna2; Plxna4* double mutants. *Plxna4* is not absolutely required for SAC lamination (Matsuoka et al., 2011a), but plexin receptor redundancy could mask PlexA4 contributions to this process. SAC lamination defects in *Plxna2^-/-^; Plxna4^-/-^* double mutants were no more severe than in *Plxna2^-/-^* single mutants (Fig. S8A). Unlike *Sema6a^-/-^* retinas, the RBC termination zone in *Plxna2; Plxna4* double mutants was maintained; as expected, TH^+^ AC dendrites mis-projected into the inner IPL. Thus, the increased severity SAC lamination defects in *Sema6a^-/-^* retinas is due to plexin-independent roles of *Sema6a*. Since loss of *Plxna4* does not exacerbate the *Plxna2^-/-^* SAC crossover phenotype, and since Sema6A is not strongly expressed in SACs when their lamination is initiated, nor is it required in SACs for their proper lamination, another cell type must influence lamination of the SAC scaffolds through a Sema6A-dependent, plexin-independent mechanism.

Similar to *Sema6a^-/-^* retinas, Syt2^+^ OFF type 2 CBC and HCN4^+^ OFF type 3a CBC axon termination domains, and also the VGlut1^+^ BC axon termination pattern were disrupted in the *Plxna2^-/-^; Plxna4^-/-^* mutants (Fig. S8B-D). Yet, the organization of these neurites was normal in *Plxna2* or *Plxna4* single mutants (*data not shown*), supporting plexin receptor redundancy in guiding the termination of BC axons. Interestingly, RBC and type 6 CBC (weak, diffuse Syt2^+^ puncta denoted by arrowheads in Fig. S8B) axon termination and VGlut3^+^ AC dendritic lamination patterns were preserved in the *Plxna2*; *Plxna4* double mutants (Fig. S8A-C), implicating plexin-independent roles for *Sema6a* in their IPL stratification.

*Plxna2^-/-^; Plxna4^-/-^*mutant retinas also exhibit mild RGC mislocalization in retinal cross sections (Fig. S8D). RGC displacement was comparable between *Plxna2^-/-^; Plxna4^-/-^* and *Plxna2^-/-^; Plxna4^+/-^* mutants, whereas RGC localization was not impacted in *Plxna2^+/-^; Plxna4^-/-^*mutants (Fig. S8D). Since RGC displacement was comparable between the *Plxna2^-/-^; Plxna4^-/-^* double mutants and *Plxna2^-/-^* single mutants, we quantified RGC displacement in *Plxna2^-/-^* retinas. In juvenile *Plxna2^-/-^* retinas, RGC count was unchanged and only 6% of RGCs were displaced (Fig. S8E-G). These results indicate that RGC displacement and loss in *Sema6a^-/-^*retinas is largely plexin-independent.

Together, our examination of various inner retina neuron projection patterns in contexts where SAC circuits are dramatically (*Sema6a^-/-^*) or partially (*Plxna2^-/-^and Plxna2^-/-^; Plxna4^-/-^*) disrupted reveals unique mechanisms underlying proper targeting of IPL neurites. For some cell types, including VGlut3^+^ ACs, RBCs, and ON type 6 CBCs, PlexA receptors are not required for neurite targeting. They are similarly not required for SAC cell body positioning, and PlexA2 is only partially required for RGC localization, suggesting that these phenotypes in *Sema6a^-/-^* mutants arise through PlexA-independent Sema6A signaling via unidentified receptor(s), or as a secondary consequence of the apicobasal disruption of the SAC scaffolds, which remain mostly intact in *Plxna* mutants. For other cell types, including SACs, OFF type 3a, and HCN4 CBCs, Sema6A/PlexA interactions are partially required for neurite targeting, as evidenced by partial disruption of their layer-specific terminations in both *Sema6a^-/-^*and *Plxna2^-/-^; Plxna4^-/-^* mutant backgrounds.

### Sema6A is Expressed by ON and ON-OFF Direction-Selective Ganglion Cells

Sema6A in SACs is dispensable for SAC lamination (Fig. 1) and Sema6A is robustly and broadly expressed by RGCs (Fig. 3 and Fig. S5). Subsets of DSGCs have immature dendritic arbors in the IPL at P1 poised to interact with SAC dendrites as they segregate (Peng et al., 2017). Therefore, we next examined Sema6A expression in RGCs to identify candidate RGC subtypes that might influence SAC scaffold elaboration.

Our previous analysis of Sema6A using anti-Sema6A immunolabeling showed clear expression of Sema6A in oDSGCs, but did not support Sema6A expression in ooDSGCs (Sun et al., 2015). Consistently, HA-Sema6A is expressed in P10 Spig1-GFP^+^ and Hoxd10-GFP^+^ oDSGCs (Fig. 6A) and their retinorecipient midbrain nuclei, including the medial terminal nucleus (MTN), nucleus of the optic tract (NOT), and dorsal terminal nucleus (DTN) of the accessory optic system (AOS) (Fig. 6B; Dhande et al., 2013; Sun et al., 2015; Yonehara et al., 2008). The superior signal:noise ratio of HA-Sema6A immunolabeling allowed us to also detect Sema6A expressed in Hb9-GFP^+^ ventral motion-preferring and Drd4-GFP^+^ nasal motion-preferring ooDSGCs (Fig. 6C), both of which project to the shell of the dorsal lateral geniculate nucleus (dLGN) and the outer shell of the superior colliculus (SC) (Zhang et al., 2017). Robust expression of HA-Sema6A was observed in the dLGN shell and the outer SC in P10 *Sema6a^HA-F/HA-F^* mice (Fig. 6D). To confirm that RGC axons are the source of HA-Sema6A in these image-forming retinorecipient targets, we performed enucleation experiments. HA-Sema6A in the dLGN shell and outer SC disappeared by one week following unilateral enucleation (Fig. S9A,B). HA-Sema6A was not expressed by RGCs projecting to non-image forming retinorecipient nuclei (Fig. S9C). Taken together, these results show that both DSGC subtypes synaptically connected to SACs, oDSGCs and ooDSGCs, express Sema6A, raising the possibility that DSGC-derived Sema6A underlies SAC circuit elaboration.

**Fig. 6.**
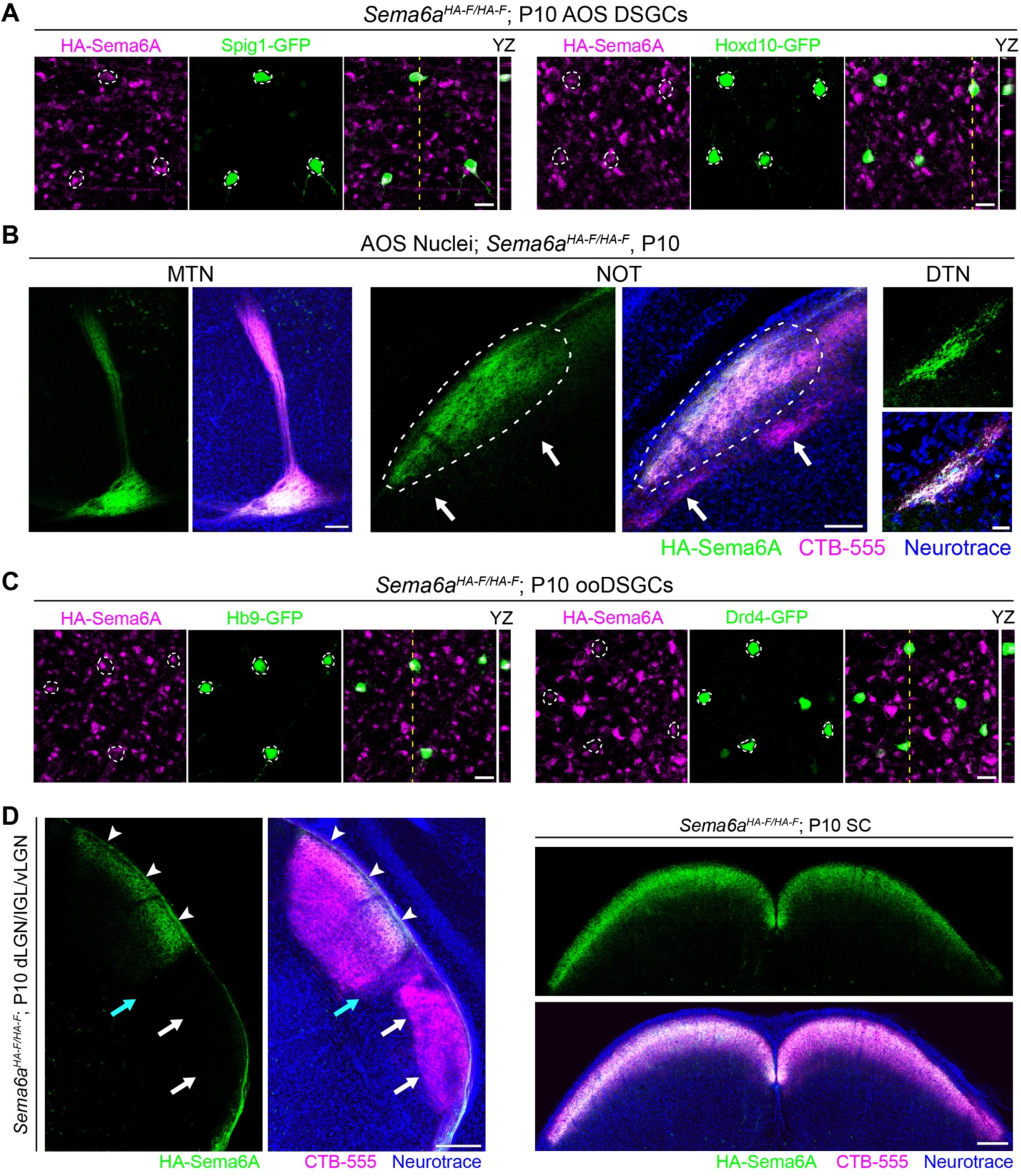
Sema6A is expressed by On and On-OFF DSGCs. **(A)**P10 Spig1-GFP^+^ oDSGCs (left) and Hoxd10-GFP^+^ DSGCs (right) in *Sema6a^HA-F/HA-F^* retinas express HA-Sema6A (magenta). Scale bar, 20µm. **(B)** HA-Sema6A (green) in P10 AOS midbrain nuclei, including the MTN, NOT (white oblong dashed line), and DTN. HA-Sema6A is not detected in the olivary pretectal nucleus (white arrows), demonstrating HA-Sema6A specificity in subsets of retinorecipient nuclei. Scale bar MTN and NOT, 100µm. Scale bar DTN, 20µm. **(C)** P10 HB9-GFP^+^ and Drd4-GFP^+^ ooDSGCs that project to image-forming retinorecipient midbrain nuclei express HA- Sema6A in *Sema6a^HA-F/HA-F^*. Scale bar, 20µm. **(D)** HA-Sema6A in the shell of the image-forming dLGN (white arrowheads), a known target of ooDSGCs, at P10. HA-Sema6A is not expressed in the dLGN core, inner geniculate leaflet (cyan arrow), or ventral LGN (white arrows). HA- Sema6A is in superficial ooDSGC target layers of the image-forming superior colliculus (right panels). Scale bars, 200µm. CTB-555 was binocularly injected at P8, labeling retinorecipient midbrain nuclei in **(B)** and **(D)**. Yellow dashed lines in **(A)** and **(C):** the region highlighted in YZ orthogonal images (right).

### Sema6A in RGCs Influences ON-SAC Plexus Elaboration and RGC Migration

*Sema6A* mRNA is expressed by postmitotic RGCs in the inner neuroblastic layer at E14.5 (Fig. S10A), prior to SAC subtype specification and IPL formation (Peng et al., 2020; Ray et al., 2018), positioning RGC-derived Sema6A to influence early retinal patterning events. Therefore, we used *VGlut2^Cre^* to conditionally remove *Sema6a* from embryonic, newly postmitotic RGCs (Wilkinson et al., 2021). HA-Sema6A is completely absent in RGC cell bodies, dendrites, and axons coursing through the optic nerve in P28 *VGlut2^Cre^*; *Sema6a^HA-F/-^* retinas (Fig. S10B). We next examined SAC plexus elaboration and RGC distribution *en face*. ON-SAC dendritic plexus organization is compromised in RGC cKO retinas, with the ON plexus exhibiting large gaps reminiscent of ON-SAC plexus disorganization in *Sema6a^LacZ^*, *Plxna2,* and *Plxna2^ΔRasGAP^* mutant retinas (Fig. 7A,B, Fig. S3D; Fig. 4E, arrowheads; Sun et al., 2013). We measured the largest diameter of these gaps and found that *VGlut2^Cre^* RGC cKO retinas had ∼150 20-29 µm diameter gaps per mm^2^, while controls had only 15 (Fig. 7B). Control ON-SAC plexuses rarely exhibited gaps >30 µm, while RGC cKO retinas had ∼50 gaps 30-39 µm across and ∼35 gaps > 40 µm per mm^2^. Many RBPMS^+^ RGCs were observed within these gaps, often clumped together with other RBPMS^-^ cells (Fig. 7A). Though the OFF-SAC plexus was not altered by *Sema6a* LOF in RGCs, many RBPMS^+^ RGCs were displaced within the OFF-SAC IPL layer (Fig. S10C). When we quantified displaced RGCs in *VGlut2^Cre^*cKO retinas, only 12% were displaced, ∼half of what we observe in *Sema6a^-/-^*mutants (Fig. 7C, 5G and 5I). The total number of RGCs in *VGlut2^Cre^* cKO retinas was not impacted (Fig. 7D), indicating RGC-derived Sema6A is not required for RGC survival (see Fig. 5F).

**Fig. 7.**
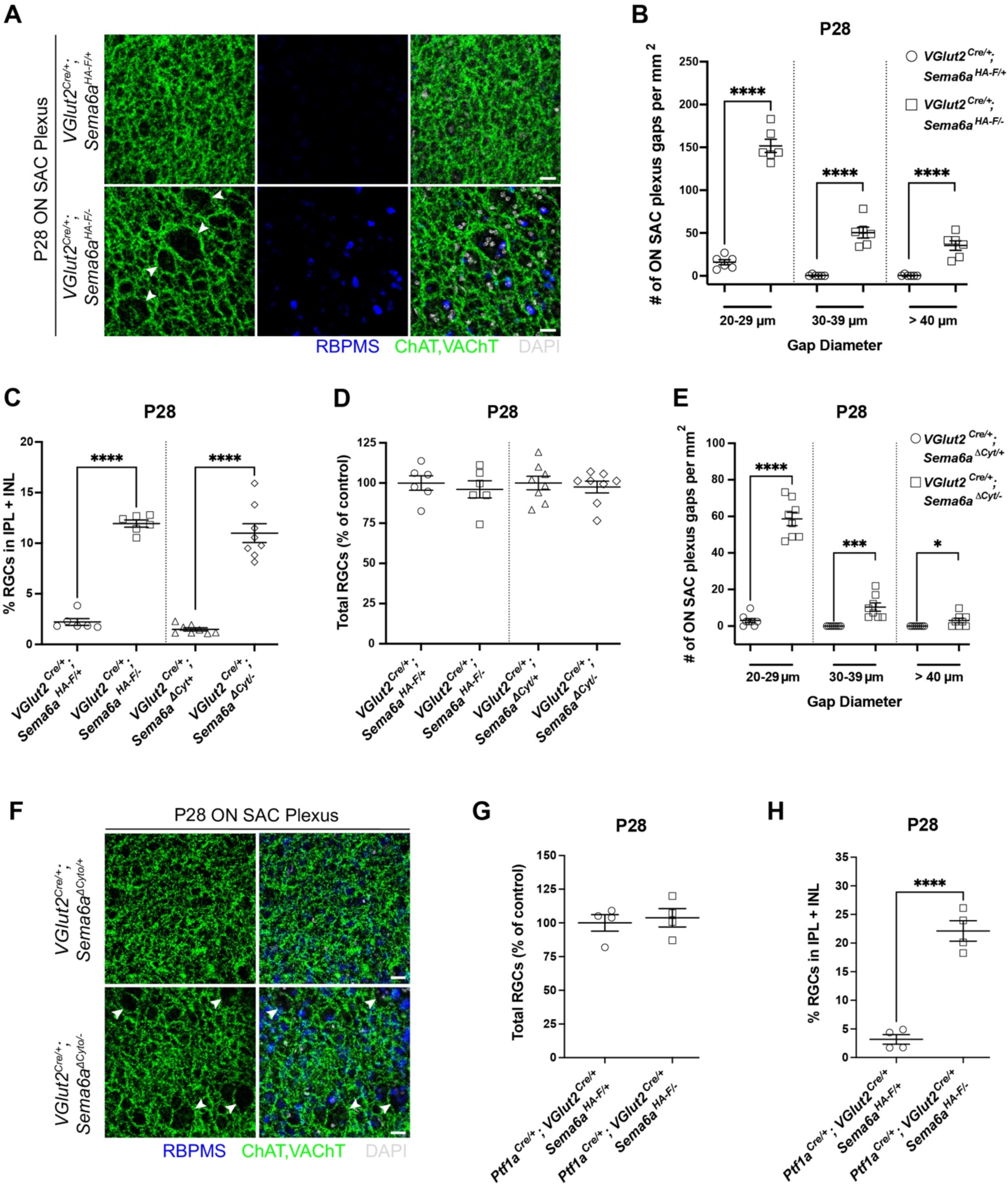
Sema6A is cell-autonomously required in RGCs for their migration and for ON- SAC plexus organization. **(A)**Large gaps appear in the P28 ON-SAC dendritic plexus in *VGlut2^Cre^; Sema6a^HA-F/-^* cKO retinas (white arrowheads). Displaced RBPMS^+^ RGCs (bottom middle) are often clustered with other DAPI^+^ cell bodies in ON-SAC plexus gaps (bottom right). **(B)** Quantification of P28 ON-SAC plexus gaps of varying diameter (in µm) per mm^2^ in *Sema6a* RGC-cKO retinas. **(C)** Quantification of IPL+INL displaced RGCs at P28. **(D)** Normalized quantification of total RGCs in RGC-cKO (*Vglut2^Cre^*) and RGC-specific Sema6A cytoplasmic deletion (*Sema6a ^ΔCyt^*) retinas at P28. **(E)** Quantification of P28 ON-SAC plexus gaps of varying diameter (in µm) per mm^2^ in RGC *Sema6a ^ΔCyt^*-cKO retinas. **(F)** Large gaps in the P28 ON-SAC dendritic plexus of *VGlut2^Cre^; Sema6a ^ΔCyt/-^* cKO retinas (white arrowheads). Quantification of total RGCs **(G)** and displaced RGCs **(H)** in pan-amacrine cell (*Ptf1a^Cre^*) and pan-RGC (*VGlut2^Cre^*) double cKO retinas at P28. Scale bars, 50µm.

To ask if Sema6A “reverse” signaling in RGCs influences SAC circuit elaboration, as it does oDSGC targeting to the MTN (Sun et al., 2015), we conditionally removed the cytoplasmic domain of Sema6A from RGCs using *VGlut2^Cre^* and *Sema6A^ΔCyt^*. Large gaps were detected in the ON-SAC plexus in *VGlut2^Cre/+^*; *Sema6A^ΔCyt/-^* retinas, but the phenotype was less severe than in *VGlut2^Cre/+^; Sema6a^HA-F/-^* cKO retinas (Fig. 7E,F compared to Fig. 7A,B). However, RBPMS^+^ RGCs were comparably displaced in *VGlut2^Cre/+^*; *Sema6A^ΔCyt/+^* and full-length cKO retinas (Fig. 7A,C,F; Fig. S10C,D). The total number of RGCs was unchanged in both full-length *Sema6a* and *Sema6A^ΔCyt^* cKO backgrounds (Fig. 7D). Therefore, Sema6A reverse signaling within RGCs contributes to RGC settling in the developing inner retina and is required for ON-SAC dendritic plexus organization, a process that might be influenced by displaced cells, including RGCs, within the plexus gaps. However, loss of reverse Sema6A signaling in *Sema6A^ΔCyt^*cKO retinas does not fully account for the ON-SAC plexus gap phenotype in full-length RGC cKO retinas, indicating that ON-SAC plexus organization is impacted by both Sema6A forward and reverse signaling events.

Only 12% of RGCs are displaced in *VGlut2^Cre^*cKO retinas, yet 25% of total RGCs are displaced in *Sema6a^-/-^* mutants, so we investigated whether another cell type is partially responsible for RGC displacement in *Sema6a*^-/-^ mutants. We examined RGC migration in P28 retinas where *Sema6a* was conditionally removed from all postmitotic ACs (Fujitani et al., 2006; *Ptf1a^Cre/+^; Sema6a^HA-F/-^*), all embryonic SACs (*Megf10^Cre/+^; Sema6a^HA-F/-^*), or in retinas where the cytoplasmic domain of Sema6A was removed from all embryonic SACs (*Megf10^Cre/+^*; *Sema6a^ΔCyt/-^*). In each of these cKO backgrounds only ∼7% of RGCs were displaced (Fig. S10E), the same fraction as in *Sema6a^+/-^* heterozygous retinas (Fig. 5G), indicating that RGC displacement in the AC cKO backgrounds stems from haploinsufficiency of the null allele and not from these genetic perturbations. Total RGC numbers were also not altered by loss of Sema6A in ACs (Fig. S10F).

Next, we generated *VGlut2^Cre^*and *Ptf1a^Cre^* double transgenic mice to conditionally remove *Sema6a* from both embryonic RGCs and ACs. Total RGC number was not impacted in *Ptf1a^Cre^; VGlut2^Cre^* double cKO retinas, but ∼22% of RGCs were displaced (Fig. 7G-H). These data suggest that Sema6A in RGCs primarily regulates RGC settling behavior through both forward and reverse signaling events. Loss of Sema6A in ACs alone was not sufficient to displace RGCs, suggesting ACs are not the source of Sema6A ligand underlying RGC positioning; but in the absence of Sema6A in RGCs (*VGlut2^Cre^* cKO), AC-derived Sema6A partially compensates to preserve ∼half of Sema6A-dependent RGC settling in the GCL. However, neither RGC- nor AC-derived Sema6A contributes to the progressive loss of RGCs in aging retinas (Fig. 7G, Fig. 5F,H), indicating that loss of Sema6A elsewhere along the visual pathway is responsible for the 26% loss of RGCs in adult *Sema6a^-/-^* retinas.

### Sema6A in RGCs Promotes SAC Dendrite Stratification

Sema6A in RGCs contributes to overall ON-SAC plexus organization; does it also regulate SAC IPL lamination? We first assessed SAC lamination in P10 *VGlut2^Cre/+^; Sema6a^HA-F/-^*retinas, which exhibit a select loss of HA-Sema6A in RGC cell bodies and axons in the nerve fiber layer (NFL) (Fig. S11A). We detected crossovers between the OFF- and ON-SAC dendrite plexuses (Fig. S11A-C), but similar to the RGC displacement phenotype of mature *VGlut2^Cre^* RGC cKO retinas, the expressivity of this crossover phenotype was not as strong as in *Sema6a^-/-^*retinas. We next examined older, P28, *VGlut2^Cre^* cKO retinas and similar to P10, HA-Sema6A protein was absent in the GCL and NFL; infrequent crossovers between OFF- and ON-SAC dendrites were detected, and the SAC strata were predominantly segregated (Fig. 8A,B).

**Fig. 8.**
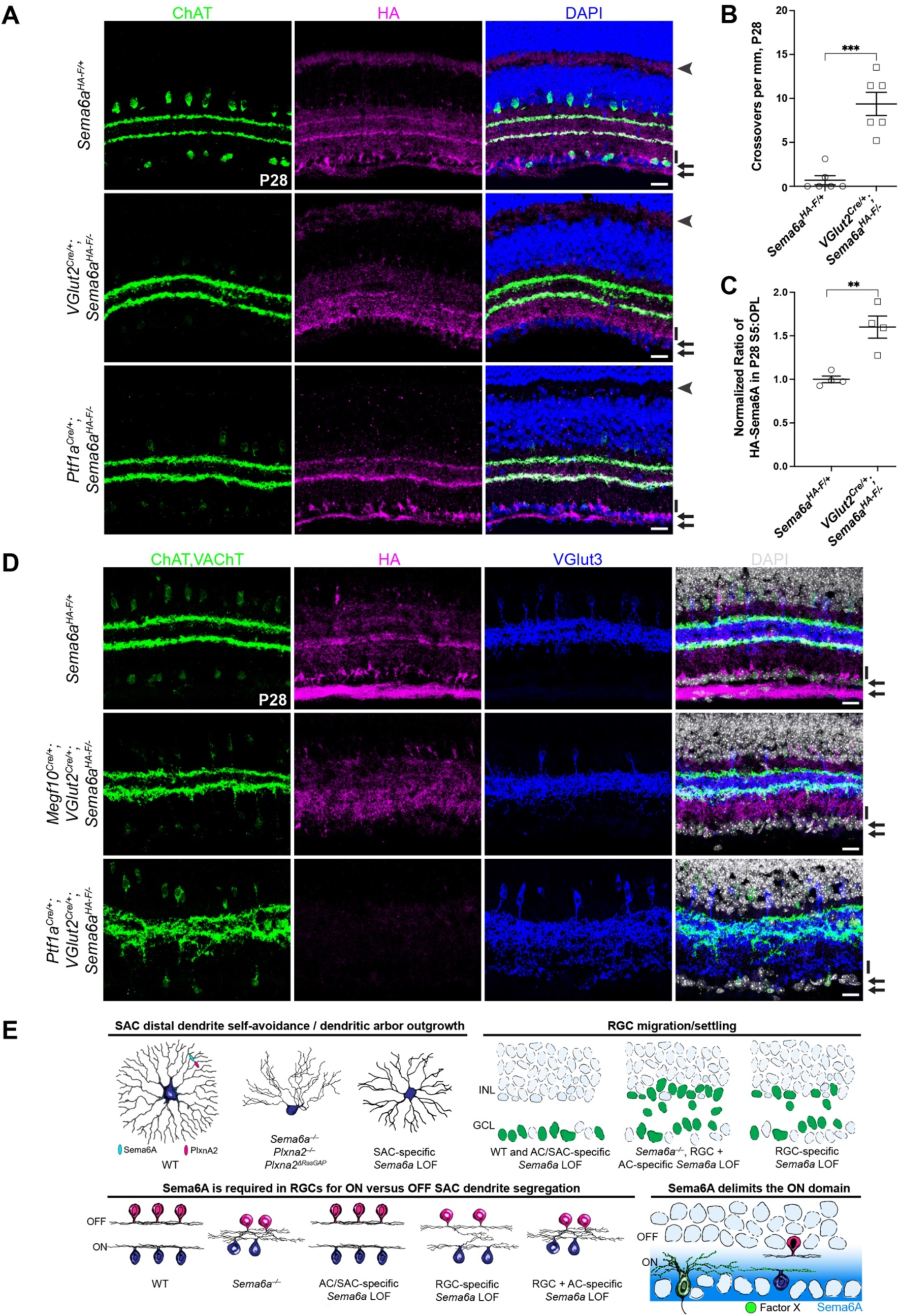
RGC-derived Sema6A segregates SAC dendrites and defines the ON domain of the IPL. **(A)**P28 retinal cross sections labeled with anti-ChAT (SACs), anti-HA (Sema6A), and DAPI. Arrowheads, right, indicate HA-Sema6A expression in the OPL; *Ptf1a^Cre^* is also expressed in horizontal cells (Fujitani et al., 2006). HA-Sema6A is not detected in the OPL of *Ptf1a^Cre^* cKO retinas. Arrows (right) indicate HA-Sema6A expression in the GCL (upper) and NFL (lower). Black bars (right) denote elevated HA-Sema6A in S5 of *VGlut2^Cre^* cKO retinas. **(B)** Quantification of the number of crossovers per mm in the IPL of P28 *VGlut2^Cre^*cKO retinas. **(C)** Quantification of HA-Sema6A in S5 of the IPL relative to HA-Sema6A in the OPL at P28. **(D)** P28 retinal cross sections labeled with anti-ChAT (SACs), anti-HA (Sema6A), anti-VGlut3 (VGlut3^+^ ACs), and DAPI. Arrows (right) indicate HA-Sema6A in the GCL (upper) and NFL (lower). Black bars (right) indicate elevated HA-Sema6A levels in S5 of *VGlut2^Cre^*cKO retinas (see **(A)**). **(E)** Model depicting roles for Sema6A in SAC and RGC development. Top left: isoneuronal Sema6A-PlexA2 signaling in SACs promotes distal dendrite self-avoidance and dendritic arbor outgrowth. Top right: RGC-derived Sema6A promotes basal RGC settling in the GCL. Bottom left: Sema6A in RGCs promotes SAC segregation; AC-derived Sema6A compensates for loss of RGC-Sema6A to maintain SAC arbor segregation in RGC-specific *Sema6a* cKO. Bottom right: proposed gradient of Sema6A expression defined by basally-biased RGC dendrites in the nascent IPL repels (pushes) SAC dendritic arbors away from the GCL, while an unknown factor X mutually expressed by ON-SACs and ON arbors of ooDSGCs attracts (pulls) ON-SAC arbors to position them basally in the IPL relative to OFF-SAC arbors. Scale bars, 20µm.

Interestingly, at both P10 (Fig. S11D) and P28 (Fig. 8C) we observed a significant increase in HA-Sema6A in S5 of the *VGlut2^Cre^* cKO IPL. Both neonatal non-SAC ACs and RGCs express Sema6A at P1 (Fig. S5C), consistent with their ability to impact SAC subtype dendrite segregation, which is complete by P1 (Ray et al., 2018). However, we found that *Sema6a* cKO from all newly born postmitotic ACs using *Ptf1a^Cre^* did not alter SAC dendrite lamination (Fig. 8A, bottom), suggesting that ACs are not the primary source of Sema6A that normally segregates SAC dendrites. Removing *Sema6a* from all ACs (*Ptf1a^Cre^*) highlights the robust expression of Sema6A in DSGC dendrites projecting to the ChAT^+^ DS circuits (Fig. 8A, bottom).

Increased S5 Sema6A expression in *VGlut2^Cre^*cKO retinas suggests that non-SAC ACs with neurites in S5 upregulate Sema6A in a compensatory fashion, preserving inner retina patterning and supporting our observation that ACs partially maintain RGC positioning in RGC Sema6A cKO retinas (Fig. 7). To ask if AC-derived Sema6A in *VGlut2^Cre^* cKO retinas occludes detection of the SAC S2-S4 fusion phenotype observed in *Sema6a^-/-^* retinas, we first removed *Sema6a* from both embryonic SACs and RGCs using *Megf10^Cre^*and *VGlut2^Cre^*, respectively, and analyzed SAC segregation. We found that OFF- and ON-SAC layers were mostly segregated with only occasional crossovers similar to *VGlut2^Cre^* cKO retinas, indicating that non-SAC AC-derived Sema6A is capable of promoting SAC strata segregation (Fig. 8D, middle).

We next conditionally removed *Sema6a* from all ACs and RGCs using *Ptf1a^Cre^* and *VGlut2^Cre^*. HA-Sema6A protein was entirely lost in the inner retina in this double cKO background, and OFF- and ON-SAC dendrites were essentially fused, similar to *Sema6a^-/-^*retinas (Fig. 8D, bottom). Additionally, while VGlut3^+^ AC neurites are properly laminated in each of the single cKO backgrounds we examined, including *Ptf1a^Cre^*(Fig. S11E), *VGlut2^Cre^* (data not shown) and also in *Megf10^Cre^; VGlut2^Cre^* cKO retinas (Fig. 8D, middle), they misproject in *Ptf1a^Cre^*; *VGlut2^Cre^*double cKO retinas – where SAC dendrites are fused – comparable to *Sema6a^-/-^*retinas (Fig. 5B). This demonstrates that for at least one cell type, VGlut3^+^ ACs, disruptions in IPL neurite lamination are a secondary consequence of the loss of discrete SAC scaffolds, rather than a direct consequence of aberrant Sema6A signaling within VGlut3^+^ neurons.

Together, these results show that: (1) ON-SAC plexus organization is compromised following loss of Sema6A in RGCs; (2) RGCs and other non-RBPMS^+^ cells are displaced within ON-SAC plexus gaps when Sema6A is removed from RGCs, which likely influences ON-SAC plexus disorganization; and (3) RGC-derived Sema6A is required for SAC dendrite stratification. Therefore, SACs rely on Sema6A expressed in RGCs for their dendritric arbor development in both planes – apicobasal and *en face* – prior to SAC scaffold-dependent regulation of apicobasal RGC dendrite lamination.

## DISCUSSION

SAC dendritic arbors carve out the nascent IPL during late embryogenesis, before the arrival of their synaptic partners, and they segregate into distinct OFF and ON layers in the IPL by P1 (Ray et al, 2018). These discrete dendritic plexuses instruct DS circuit formation by serving as a scaffold upon which DSGC dendrites and BC axons refine and form synaptic connections (Stacy and Wong, 2003; Peng et al., 2017; Duan et al. 2018). In the present study, we generated an epitope-tagged *Sema6a* cKO allele that permitted an in-depth investigation of Sema6A-mediated signaling events impacting SAC scaffold elaboration during retinal development. We predicted that loss of *Sema6a* in SACs would phenocopy the IPL lamination deficits of *Sema6a^LacZ^* retinas and confirm our hypothesis that Sema6A in ON-SACs repels OFF-SACs (Sun et al., 2103). We were surprised to discover instead that RGC-derived Sema6A affects SAC scaffold segregation, indicating that these earliest born retinal neurons help define key features of OFF versus ON visual circuits prior to layer-specific targeting and refinement of their own synaptic connections within the IPL.

### Sema6A is Expressed in All SACs and Promotes Distal Dendrite Self-Avoidance

We selectively removed *Sema6a* from all SACs and found that Sema6A within SACs is dispensable for SAC lamination, a critical first step in DS circuit formation (Peng et al., 2017; Duan et al., 2018). By conditionally removing *Sema6a* in isolated SACs we confirmed that isoneuronal Sema6A signaling promotes SAC distal dendrite self-avoidance and outgrowth. We also found that isoneuronal forward Sema6A signaling through PlexA2, involving the PlexA2 RasGAP domain, is required for OFF- and ON-SAC distal dendrite self-avoidance, consistent with our observation that OFF-SAC dendritic outgrowth is diminished in *Plxna2^-/-^* retinas (Sun et al., 2013). Examination of isolated SAC morphology in *Sema6a^-/-^*retinas revealed that *Sema6a* is also required for OFF-SAC outgrowth and, to a lesser degree, dendritic arbor symmetry. Furthermore, the superior specificity and signal of HA-Sema6A immunolabeling, coupled with SAC scRNA-seq and characterization of *Sema6a^CreER^* reporter activity, facilitated identification of Sema6A expression in OFF SACs, in addition to ON SACs. These results demonstrate that forward isoneuronal Sema6A-PlexA2 signaling in all SACs promotes distal dendrite self-avoidance, which is partially responsible for SAC dendritic arbor outgrowth and radial morphology. While we favor the idea that Sema6A signals through PlexA2 *in trans* in SACs, we cannot rule out contributions of *in cis* signaling. These results underscore the complementarity between distinct dendritic self-avoidance mechanisms, whereby protocadherin signaling plays a critical role in SAC primary dendrite self-avoidance (Lefebvre et al., 2012) and Sema6A-PlexA2 signaling influences distal dendrite self-avoidance (Sun et al., 2013; this study).

The ON-SAC outgrowth defect was not as severe in *Sema6a* cKO SACs compared to *Sema6a^-/-^*ON-SACs. The timing of *Chat^CreER^* expression may contribute to this discrepancy, but it is likely that atypical repulsive interactions between SACs and displaced retinal neurons/neurites underlies the severe ON-SAC outgrowth defect of *Sema6a^-/-^* retinas. In SAC-specific cKO retinas, ON-SAC radial symmetry and overall plexus organization are preserved. In genetic backgrounds where ON-SAC radial symmetry is disrupted, including *Sema6a^-/-^*, *Sema6a^LacZ/LacZ^*, *Plxna2*^-/-^, and *Plxna2^ΔRasGap^* mutants, large gaps interrupt the ON-SAC plexus. These gaps correlate with regions where ON-SACs fail to elaborate their dendritic arbors (Sun et al., 2013) and often contain displaced RGCs and/or cells of unknown identity. Gaps are also observed in RGC cKO retinas, where RGC positioning is disrupted, suggesting that repulsive interactions from displaced RGCs within the IPL disrupt the ON-SAC plexus. We find that SAC dendritic arbors in primary retinal cultures collapse if they contact other cell types (*data not shown*), supporting the idea that SAC dendritic arbor symmetry is sensitive to non-SAC interactions.

Interestingly, many RGCs in *VGlut2^Cre^*cKO retinas are also displaced within the OFF IPL layers and the basal-most AC zone of the INL, and yet overall OFF-SAC plexus organization is not altered. ON-SAC plexus disorganization may be a secondary consequence of exposure to repulsive signaling distinct from Sema6A that ON-SACs are selectively competent to respond to but would otherwise be isolated from under normal conditions. Alternatively, displacement of RGC subtypes into only the ON layers of the IPL may preclude OFF-SAC exposure to repulsive cues only ON-SACs engage. Still, it is possible that loss of Sema6A in a subset of RGCs removes a positive signaling pathway that promotes ON-SAC dendrite outgrowth and symmetry *en face*. Further examination of RGC subtypes displaced in *Sema6a^-/-^*retinas may shed light on the mechanisms underlying preferential ON-SAC dendritic arbor disruption in various mutants that impact Sema6A-PlexA2 signaling.

### Sema6A is Required for Precise Positioning of RGCs

Sema6A-PlexA2 signaling has been implicated in interkinetic nuclear migration of retinal progenitor cells (Belle et al., 2016). We show here that Sema6A, and to a lesser degree PlexA2, also contributes to migration of postmitotic RGCs. Previous studies suggest that RGCs migrate in a bipolar somal translocation fashion (Hinds and Hinds, 1974; Morest, 1970), yet *in vivo* light-sheet imaging of embryonic zebrafish retina identified two phases of RGC migration: 1) rapid, directional bipolar somal translocation into the basal retina; and 2) a slower phase of RGC “fine-positioning” that is likely related to RGC subtype mosaic spacing (Icha et al., 2016). When basal RGC migration was impaired in this zebrafish study, RGCs stalled near the apical surface of the retina. Here, basal RGC migration through the outer neuroblastic layer was entirely preserved. However, 26% of RGCs failed to settle in the GCL of *Sema6a^-/-^*retinas, implicating defective fine-positioning of RGCs in the absence of Sema6A.

Conditional removal of either full-length *Sema6a* or its cytoplasmic domain in postmitotic RGCs recapitulated ∼1/2 of the RGC displacement phenotype observed in *Sema6a^-/-^*mutants, indicating that reverse signaling in RGCs is required for GCL settling. However, the full expressivity of the *Sema6a^-/-^* RGC displacement phenotype was not achieved until Sema6A was removed from postmitotic RGCs and ACs in *Ptf1a^Cre^*; *VGlut2^Cre^* double cKO retinas. Reverse Sema6A signaling in RGCs contributes to RGC settling, while AC-derived Sema6A normally does not. *Sema6a* cKO in ACs and RGCs together fully recapitulates the *Sema6a^-/-^* RGC settling phenotype, suggesting that while RGCs are likely the sole source of Sema6A in this process, ACs can compensate to confine RGCs to the GCL.

In migrating cerebellar granule neurons, Sema6A coordinates a switch from early tangential to later radial modes of migration (Kerjan et al., 2005), and lack of Sema6A-PlexA2 signaling specifically impairs migration of later-born cerebellar granule neurons (Renaud and Chédotal, 2014). We propose that in the developing mouse retina, Sema6A promotes fine-scale RGC settling, specifically affecting later-born RGCs to account for only partial (26%) displacement of the RGCs in *Sema6a^-/-^*retinas. Reverse Sema6A signaling was recently shown to suppress migration of H1299 lung cancer cells (Chen et al., 2019), supporting a similar role in RGCs to either suppress basal migration or to terminate exploratory fine-positioning (“settling”) migratory behavior. Termination of basal migration appears to be preserved in *Sema6a^-/-^* retinas since RGCs do not migrate beyond the GCL. Therefore, forward and reverse Sema6A signaling in later-born RGCs likely coordinately promote the switch from basal to fine settling migration and may facilitate termination of exploratory fine-positioning in some RGCs.

### Sema6A in RGCs Delimits the ON Domain of the IPL

The establishment of distinct OFF versus ON IPL domains, where neurites are brought into close apposition that depolarize in response to light decrements increments, respectively, is a key developmental event for patterning retinal circuit connectivity (Zhang et al., 2017). Since SAC subtypes are specified prior to IPL generation (Peng et al., 2020), SAC subtype specification and elaboration of the distinct OFF- versus ON-SAC dendritic scaffolds are the earliest events that pattern the IPL, setting the stage for later OFF and ON RGC, AC, and BC neurite targeting. Next, the inner retina is delineated from the outer retina, and critical roles in this early patterning event require repulsive signaling by the transmembrane proteins Sema5A and Sema5B, acting through PlexA1 and PlexA3 receptors (Matsuoka et al., 2011b). Here, we find that Sema6A expressed by postmitotic RGCs, the earliest-born retinal neuron subtype, initiates the patterning of IPL OFF versus ON circuits by instructing SAC scaffold segregation, facilitating the generation of the “ON” IPL domain.

Since we observed robust Sema6A expression in neonatal RGCs, we expected that RGC *Sema6a* cKO would recapitulate the *Sema6a^-/-^* SAC lamination defect. However, we found only a modest crossover phenotype between SAC strata in RGC cKO retinas, but we noted an apparent ∼2.5-fold increase in Sema6A expression in non-SAC AC neurites in the inner ON IPL layers when Sema6A was selectively removed from RGCs. This upregulation of Sema6A within the ON IPL in RGC-specific cKO retinas indicates that some ACs sense Sema6A expression and compensate for insufficient levels, ultimately maintaining overall OFF versus ON inner retinal patterning. This compensatory mechanism highlights the importance of basal IPL Sema6A expression for ON- versus OFF-SAC dendritic arbor segregation, IPL patterning, and matching pre- and postsynaptic partners within distinct OFF and ON circuits.

There is a greater abundance of Sema6A in ON (S4) compared to OFF (S2) DSGC dendritic arbors, as revealed by AC-specific *Sema6a* cKO. We attribute this asymmetry to the predominance of oDSGC dendritic arbors in S4 of the IPL, since oDSGCs project only a small fraction of their dendrites into S2 (Dhande et al., 2013). Although DSGC dendrites are not fixed in discrete IPL laminae by P1 (Kim et al., 2010), when SAC lamination is complete (Kay et al., 2018), they occupy the IPL in a graded fashion with an increased dendrite density adjacent to the GCL (Peng et al., 2017). Thus DSGC dendrites could establish a Sema6A gradient capable of assembling ON and OFF-SAC scaffolds. We propose these mixed ON and OFF DSGC dendrites at this early stage of retinal development first repel all SAC dendrites to set up the inner ON IPL domain. Subsequently, a push/pull mechanism may sort SAC dendrites. In this scenario, Sema6A on RGCs would push developing SAC arbors away from the GCL, while a select pull mechanism between ON-SAC dendrites and oDSGC dendrites, for example CNTN5- mediated attraction between ON-SACs and ON arbors of ooDSGCs (Peng et al., 2017), counters the repulsion of the ON-SAC dendritic arbor to fix their position closer to the GCL. Since *Cntn5* loss-of-function does not impair SAC dendrite segregation, and ooDSGCs and ON-SACs express multiple CNTN and contactin-related protein (Caspr) isoforms (Rheaume et al., 2018; data not shown), it is likely that compensatory adhesive mechanisms ensure that the ON-SAC dendritic arbor is elaborated basal to the OFF-SAC arbor. These include local signaling employing FLRT2-LPHN adhesive interactions that regulate ooDSGC dendrite targeting, and that are negatively regulated by UNC5 binding to FLRT2 (Prigge et al., 2023).

Taken together, our analysis of Sema6A signaling during retinal development, made possible by new genetic reagents, highlights a general requirement for SAC scaffolds in elaborating ON versus OFF domains within the IPL, a critical early step for functional circuit segregation in the developing retina. We observe that SAC scaffold development is in part enabled by isoneuronal Sema6A–PlexA2 signaling in both ON- and OFF-SACs. However, we also find that the early segregation of ON- versus OFF-SAC dendrites, which is required for IPL patterning of many retinal neuron type neurites, is unexpectedly regulated by Sema6A in RGCs, the earliest born neurons in the retina.

## MATERIALS AND METHODS

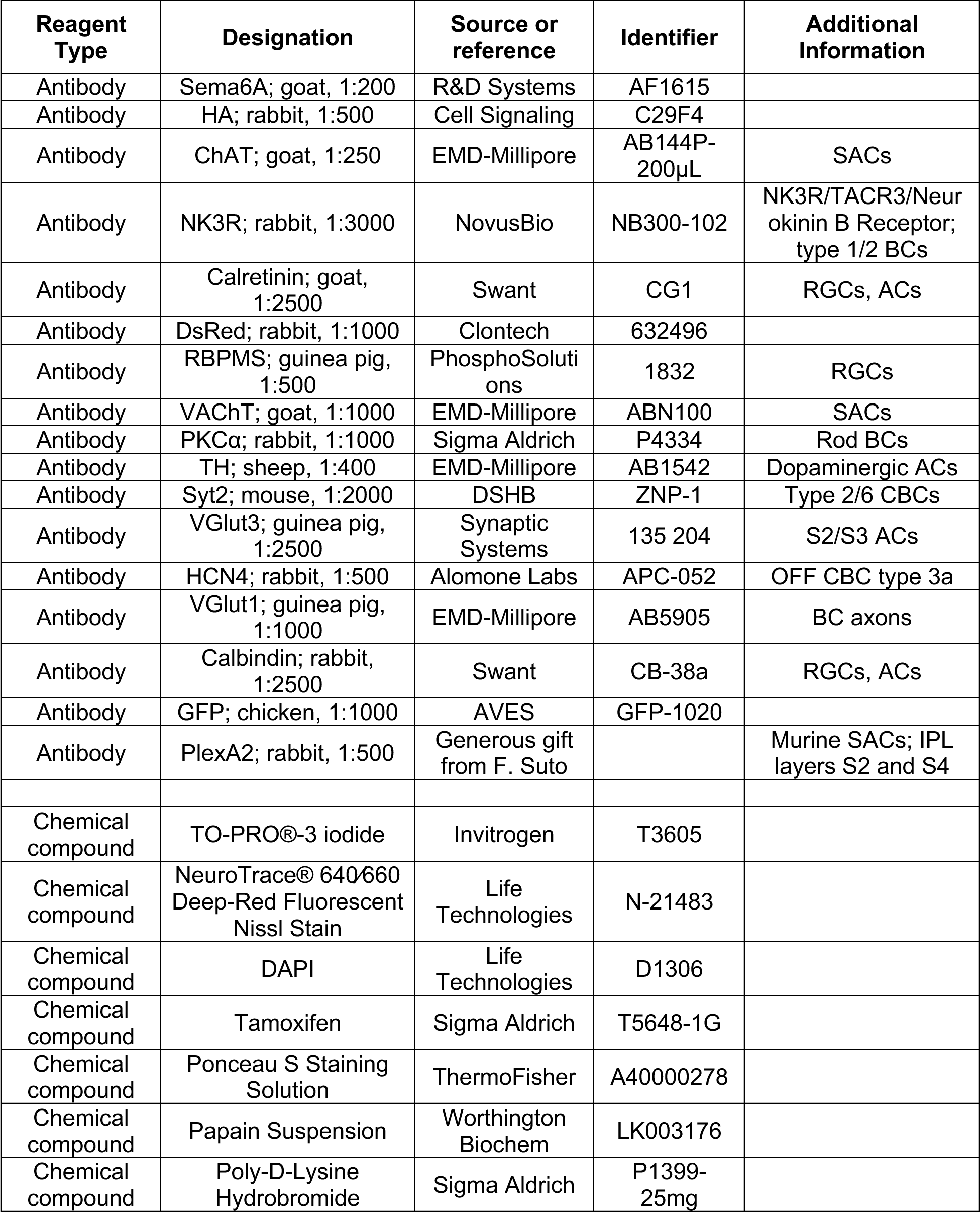
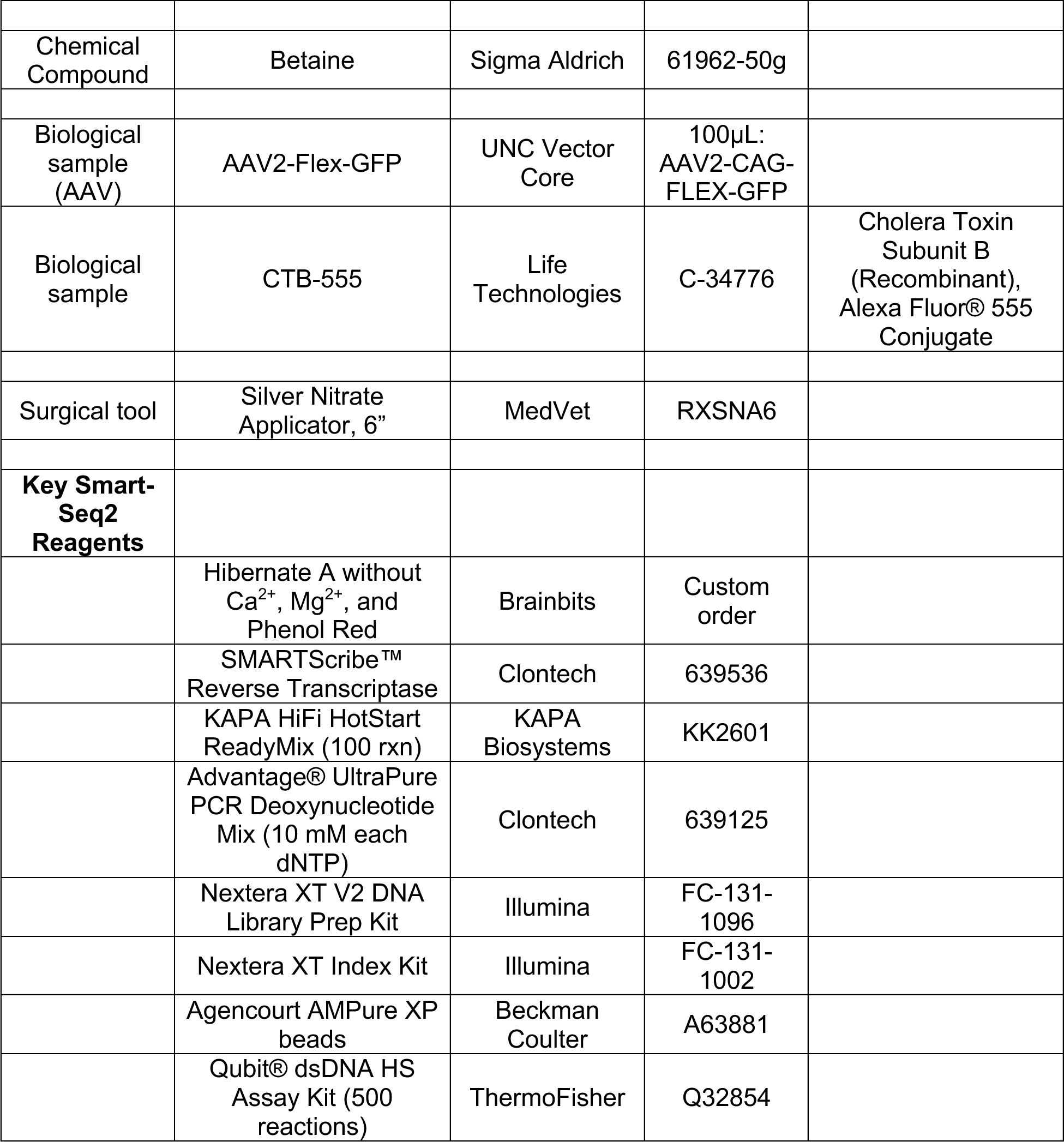
Key Resource Table.

### Animals

All animal experiments and procedures were approved by the Johns Hopkins Animal Care and Use Committee of the Johns Hopkins University. Mice (*Mus musculus*) were maintained in pathogen-free facilities at Johns Hopkins under standard housing conditions with continuous access to food and water. Histological studies per performed using postnatal day 1 (P1) to adult mice as indicated. Mice of both sexes were used, and no sexual dimorphisms were observed in any results reported here. None of the mice had noticeable health or immune status abnormalities and were not subject to prior procedures. The genotype of mice is described where appropriate. For all genetic experiments, an n of at least 4 retinas from two animals was examined, unless otherwise noted. Littermates were used as control animals, unless otherwise noted. The following mouse lines were used:

1. In *Sema6a^HA-F^*, a hemagglutinin (HA) tag is knocked in downstream of the signal peptide in exon 2 at the N-terminus of the extracellular domain of Sema6A. *LoxP* sites were inserted in introns flanking exon 3. The insertion was verified by sequencing. Recombination between the *loxP* sites introduces a frameshift and multiple premature stop codons, preventing expression of Sema6A. *Sema6a^HA-F^* was generated by the Gene Targeting & Transgenics Facility at Janelia Research Campus.
2. To generate the *Sema6a* null allele, an *Sema6a^HA-F^*male was bred with a *Sox2^Cre^* (see below) female, in which Cre recombinase is expressed in the germline. *Sema6a^HA-F^* ^/+^; Sox2^Cre/+^* (*, recombined cKO allele) offspring were then backcrossed to C57BL/6 wild type mice for five generations to 1) segregate *Sox2^Cre^*away from the Sema6a null allele, and 2) create an isogenic stock.
3. In *Sox2^Cre^* (Jackson Labs 008454, Hayashi et al., 2002), Cre recombinase is expressed under control of the mouse SRY-box containing gene 2 promoter. In *Sox2^Cre^* animals, Cre recombinase activity is detected in the epiblast cells as early as embryonic day 6.5; Cre expression is present in the female germline.
4. In *Six3^Cre^* (Jackson Labs, 019755, Furuta et al., 2000) Cre recombinase is expressed under control of the sine oculis-related homeobox 3 (Six3) promoter. Six3 is a transcription factor that is required for the development of the eye. *Six3^Cr^*^e^ is expressed ubiquitously in the central retina, but is largely absent in the peripheral retina (Ray et al., 2018).
5. In *Chat^Cre^* (Jackson Labs 006410, Rossi et al., 2011), Cre recombinase activity is weakly evident in neonatal SACs and increases over early postnatal development (Ray et al., 2018).
6. In *ROSA^LSL-TdTomato^*, also known as Ai14 (Jackson Labs, #007914, Madisen et al., 2010), the red fluorescent variant TdTomato is expressed in a tissue-specific manner depending on the cell-type specific expression of Cre recombinase.
7. In *Chat^CreER^* (Jackson Labs #008364, Rotolo et al., 2008), tamoxifen-inducible Cre recombinase is expressed in neonatal SACs in the retina and across development upon administration of tamoxifen.
8. In *Sema6a^LacZ^*, *Sema6a* gene trap mouse line (Leighton et al., 2001); determined here to be a hypomorphic allele.
9. In *Megf10^Cre^*, in Cre was inserted in the *Megf10* locus so that endogenous Megf10 expression is preserved, just downstream of the Meg10 coding sequence, with the Megf10 and Cre reading frames separated by a T2A self-cleaving peptide sequence (Peng and James et al., 2020).
10. In *ROSA^LSL-TdTomato^* (B6.Cg-*Gt(ROSA)26Sor^tm14(CAG-tdTomato)Hze^*/J, Jackson Labs #007914), a *loxP*-STOP-*loxP* cassette prevents expression of TdTomato in the absence of Cre recombinase.
11. In *Sema6a^CreER^*, a targeting construct was designed to insert the coding sequence for tamoxifen-inducible Cre recombinase fused with the mouse G525R mutant estrogen receptor 1 (ER; Schwenk et al., 1998) downstream of the stop sequence (which was removed) in exon 19 of the *Sema6a* locus. A furin cleavage RAKR sequence was placed directly downstream of the Sema6A coding sequence to separate Sema6A and the CreER fusion protein. A 2xV5 linker was added just downstream of the furin cleavage sequence to label the N-terminal fragment of the subsequent P2A self-cleaving peptide separating the Sema6A and CreER coding sequences. The *Sema6a^CreER^*allele was generated by the Gene Targeting & Transgenics Facility at Janelia Research Campus.
12. In *Plxna2^R1746A^* (Zhao et al., 2018), here referred to as *Plxna2^ΔRasGAP^*, CRISPR/Cas9 gene editing awas used to generate a point mutation that replaces a catalytically essential arginine reside with alanine, rendering *Plxna2^ΔRasGAP^* GTPase activating protein (GAP)-deficient.
13. In *Plxna2* nulls, insertion of the ITak-Neo cassette in the Plxna2 locus generates only the truncated PlexA2 signal peptide (Suto et al., 2007).
14. In *Plxna4* nulls, insertion of the PGK-neo cassette replaces exons 18 and 19, which contain the transmembrane domain, of *Plxna4*, generating a null allele (Yaron et al., 2005).
15. In *VGlut2^Cre^* (B6J.129S6(FVB)-*Slc17a6tm2(cre)Lowl*/MwarJ, Jackson Labs #028863), Cre recombinase is expressed in glutamatergic neurons under control of the vesicular glutamate transporter 2 promoter (Vong et al., 2011).
16. In *Sema6a^ΔCyt^*, exon 19 of the *Sema6a* locus, which includes the intracellular domain (ICD) coding sequence of Sema6A, was flanked by loxP sites. Cre-dependent recombination of this allele generates Sema6A lacking its ICD that can act only as a ligand. A complete description of this line is presented elsewhere (Verhagen et al., 2023).
17. In *Ptf1a^Cre^*, the Cre coding sequence replaced the coding sequence for Ptf1a at the translation initiation codon (Kawaguchi et al., 2002). The basic helix-loop-helix transcription factor *Ptf1a* is expressed in precursors of amacrine and horizontal cells from E12.5 to P3 in the murine retina (Nakhai et al., 2007).

### SAC Culture

P0-P2 *ChatCre; ROSA^LSL-TdTomato/+^*(Fig.SX) or *Sema6a^HA-F/HA-F^* (Fig.SX) retinas were dissociated using Worthington Biochem Papain supplemented with DNase I and L-Cysteine, and were cultured according to Lefebvre et al., 2012. Briefly, SACs were cultured on poly-D-lysine hydrobromide coated glass coverslips at a density of 15-20k cells per well for 8-10 days, with media refreshment every 2-3 days: one-third of the media was replaced with fresh media. Importantly, 15% of mouse cortical astrocyte-conditioned media was added to the RGC culture medium. Astrocyte-conditioned media was prepared as in Albuquerque et al., 2009. SACs in (Fig.SX) were identified by virtue of TdTomato expression. SACs in (Fig.SX) were identified based on their unique radial morphology. Density was optimized to yield cultures with isolated SACs, since SAC morphology *in vitro* is compromised by contact other cells.

### Immunohistochemistry

Mice (*Mus musculus*) were euthanized by continuous exposure to CO2 followed by cervical dislocation. Eyes were harvested and fixed with 4% PFA for 1-2 hrs at room temperature, and rinsed with PBS. For retinal cross sections, the lens was removed and eyes were immersed in 30% sucrose until sinking before embedding in Neg-50 frozen section medium (Fisher). Retinas were cryosectioned at 20µm onto Superfrost Plus slides (Fisher). Antibodies were diluted in antibody diluent (0.02% PBST plus 10% host antibody compatible serum) and were incubated with sections overnight at 4° in a humidified chamber. Sections were washed with PBS before incubating with secondary antibodies (1:500) and DAPI in antibody diluent for 1-2hrs at room temperature in a humidified chamber. Sections were further washed with PBS before mounting in Fluoro-Gel with DABCO (EMS). For whole mount retina staining, retinas were dissected and permeabilized in PBS/BSA/Triton-X permeabilization solution overnight with turning. Permeabilized retinas were incubated with primary antibodies in an antibody diluent consisting of 0.02% PBST plus 10% host antibody compatible serum with 20% DMSO for 5-7 days at room temperature with turning. Retinas were washed 5-6 times with PBS for 1hr at room temperature, with turning, before incubating in secondary antibodies (1:500) plus DAPI in antibody diluent for 1-2 days at room temperature with turning. Retinas were washed 5-6 times with PBS for 1hr at room temperature before mounting onto Superfrost Plus slides (Fisher) in Fluoro-Gel with DABCO (EMS). Broken coverslip pieces were placed at the 4 corners beneath the coverslip to prevent contact between the retinas and the coverslip. Slides were meticulously sealed with nail polish to prevent evaporation of the Fluoro-Gel.

### Image Acquisition and Analysis

Images were acquired on Zeiss LSM 710 confocal microscopes with 405, 488-515, 568, and 647 lasers, processed using Zeiss ZEN software, and analyzed using ImageJ (NIH). Section images were acquired with 20X air or 40X oil lens at the resolution of 1024×1024 pixels, a step size of 0.5-1.5 µm. ImageJ (NIH) software was used to generate maximum intensity projections. Adobe Photoshop CS6 was used for adjustments to brightness and contrast.

### Surgical Procedures

Intravitreal injections of AAVs (AAV2-Flex-GFP, UNC Vector Core) or cholera toxin B (CTB, Life Technologies) were performed as previously described (Sun et al., 2015), with animals anesthetized by isofluorane (3% in O2). A 30 gauge needle was used to make an incision at the limbus. A 33 gauge Hamilton syringe was used to slowly introduce 1µL of AAV or CTB; the needle was removed 30 seconds after the injection, to allow for dispersion of the solution into the intravitreal space. For AAV experiments, tissue was collected two weeks after injection, to allow for sufficient expression. For CTB injections before eye opening, the fused eyelid was opened using Vannas spring scissors (Fine Science Tools, 15003-08). Animals were recovered on a heated pad.

For enucleation, animals were deeply anesthetized using isofluorane (3% in O2), and enucleation was performed as in Lim et al., 2016. Briefly, tissue surrounding the eye was depressed slightly with blunt forceps, and curved surgical scissors (ROBOZ, RS-5675) were used to slightly elevate the eye from the orbit, and the optic nerve was swiftly cut and the eye removed. Afterwards, a silver nitrate applicator was gently and swiftly (∼1 sec) turned in the orbit to cauterize the tissue. Animals were recovered on a heated pad. CTB was intravitreally injected in the remaining eye at P14 prior to tissue collection at P15.

For tamoxifen-inducible expression of *ROSA^LSL-TdTomato^*using *Sema6a^CreER^*, tamoxifen was injected into the milk sac of P1-P3 pups; to examine expression at P6-P14, tamoxifen was injected subcutaneously (n = 3 per timepoint). For neonatal recombination of *Sema6a^HA-F^*in Fig. 2, tamoxifen was injected into the milk sac of P0.5 pups, after milk was evident.

### Smart-Seq2 scRNA-seq

Single cell mRNA sequencing was performed using a modified Smart-seq2 protocol that has been described in detail elsewhere (Chevee et al., 2018). Briefly, 4-6 retinas from 3-6 *Chat^Cre/+^;* ROSA^LSL*-TD-Tomato*^ mice were isolated at P1, P4, P8, P11 and P16 in ice cold HBSS, and the retinal pigmented epithelium and inner limiting membranes were removed. Retinas were digested with papain, triturated slowly 15-20 time with a P1000 pipet tip, strained into a 50 mL conical tube, and pelleted for 3 minutes at 1.3k rpm. The pellet was gently resuspended and cells were washed three times in ice cold Hibernate A media, with pelleting between the washes. Care was taken to tap the tube gently to dislodge the pellet before adding media and resuspending the pellet, to prevent cell lysis and subsequent accumulation of DNA and cellular debris, which forms as a sticky gel-like substance in the supernatant. Washed and resuspended cells in ice cold Hibernate A were subjected to FACS to isolate single cells in cell lysis buffer before proceeding with Smart-seq2 mRNA sequencing as in Chevee et al., 2018. Libraries were sequenced with the Illumina NextSeq platform at the Johns Hopkins Genetics Research Core Facility to generate 75 bp paired-end reads at mid output (150M reads).

### Primer Sequences

oligo-dT30VN:

/5Biosg/AAGCAGTGGTATCAACGCAGAGTACTTTTTTTTTTTTTTTTTTTTTTTTTTTTTTVN ISPCR: /5Biosg/AAGCAGTGGTATCAACGCAGAGT

TSO: /5Biosg/AAGCAGTGGTATCAACGCAGAGTACATrNrG+G

scRNA-seq FASTQ files were aligned to the mouse reference genome GRCm38.p6 (mm10) using HISAT2 v2.1.0 (Kim et al., 2015). SAM files were converted to BAM files using SAMtools v1.3.1. Transcript abundance was quantifed with using Cufflinks v2.2.1 (Trapnell et al., 2012). FPKM values from Cufflinks were used as input for downstream analyses in the Monocle2 framework (Trapnell at el., 2014) in RStudio v1.4.1103 or Seurat (Satija et al., 2015) in R v3.6.2, to generate violin plots. Normalized FPKM values were converted to estimated copies per cell using the Census algorithm (Qiu et al., 2017). Cells with expression of >2000 and <50,000 mRNAs were selected for further analysis. OFF and ON SAC clusters were defined by virtue of *Rnd3* and *Fezf1* expression Peng and James et al., 2020). The total number of ON and OFF SACs sequenced and analyzed at each timepoint are as follows:

### Quantification and Statistical Analysis

In Fig. 1, the germline and pan-retinal *Sema6a* cKO SAC lamination defects were too pronounced to score single crossovers between the OFF and ON strata. To overcome this we drew and measured lines in ImageJ to define the occurrence of parallel and uninterrupted ChAT^+^ SAC immunolabeling between the OFF (S2) and ON (S4) SAC strata per 320µm in 320µm x 320µm images. An n ≥ 3 animals per genotype was quantified, with three images per retina averaged, and the value of both retinas per animal averaged. For crossovers between S2 and S4 (Fig. 4B, Fig. 8B, and Fig. S8C), ChAT^+^ signal spanning both strata that was ≥ 2 µm wide was considered a *bona fide* crossover, and independent crossovers were scored as such when separated by at least 20 µm. Given this stringent scoring method, which was chosen to exclude duplicate scoring of single arbors crossing over more than once and to capture crossovers of independent SACs/SAC arbors, it is possible that crossover events were undercounted. SAC dendritic field areas were quantified in ImageJ by joining a segmented line drawn around the perimeter of the dendritic arbors, and SAC symmetry indices were calculated as in Sun et al., 2013 (n ≥12 SACs per genotype in Fig. 2B and Fig. 3C, n = 8 per genotype in Fig. 2E). To quantify gaps in the ON SAC plexus, a line was drawn within each gap as its largest diameter, and only gaps > 20 µm were considered a *bona fide* gap, versus the normal holes within the honeycomb structure of SAC arbors. When quantifying RBPMS^+^ RGC displacement, a Z-stack was taken in each quadrant of a whole mount retina flattened by 4 releasing cuts placed ∼90° apart. The image was centered as much as possible along the central to peripheral retinal axis in each quadrant. Total RGCs in register with DAPI nuclear counterstain in each 320µm x 320µm image were counted throughout the stack, and quadrant values were averaged to generate a values (% of displaced RGCs, or total RGCs) per retina. For quantification of HA-Sema6A protein in S5 of the IPL following *Sema6a* cKO in RGCs (Fig. 8C and Fig. S11D), the segmented line tool in ImageJ was used to draw a closed object around the ChAT^+^ ON SAC band in S4. This object was then moved down into S5, adjacent to the DAPI^+^ nuclear GCL, and HA-Sema6A signal was measured. A segmented line was subsequently drawn in the OPL and the HA-Sema6A signal there was measured to serve as a source of HA-Sema6A signal independent of that in the IPL for normalization. The ratio of HA- Sema6A in S5/OPL for three images per retina for two mice per genotype was averaged per experiment (n = 4 retinas from 2 mice).

All data are shown as mean ± SEM with n representing the sample number from at least four retinas from two mice, if not explicitly designated otherwise. For neuronal morphology quantifications, at least 8 neurons were quantified. Power calculations were not performed to predetermine the sample size due to our limited ability to generate many mice with complicated genetic backgrounds. Data was not excluded from any analysis. Two-tailed Student’s t-tests were used for comparisons between two groups, and one-way ANOVA with post hoc Tukey HSD was used for multi-group comparisons. Asterisks represent conventional annotations: *, <0.05; **, <0.01; ***, <0.001; ****, <0.0001.

## ACKNOWLEDGMENTS

We thank Roman Giger for *Plxna2^R1746A^* mice, Lisa Goodrich and Christopher Wright for *Ptf1a^Cre^* mice, Jeremy Kay for *Megf10^Cre^*mice, and Marieke Verhagen for assistance with *Sema6a^Cyt^* mice. We thank Caiying Guo of the Gene Targeting & Transgenics Facility at Janelia Research Campus, hhmi, for generating the *Sema6a^HA-F^* and *Sema6a^CreER^* alleles. We thank Fumikazu Suto for PlexA2 antibodies. We thank Genevieve Stein-O’Brien and Brian Clark for technical assistance and helpful discussions related to scRNAseq. We thank Hao Zhang of the Bloomberg Flow Cytometry and Immunology Core at Johns Hopkins for his excellent assistance FACS isolating SACs. We also thank Dr. Matt Brown for his excellent technical assistance, and all Kolodkin laboratory members for assistance and discussions throughout the course of this project.

## AUTHOR CONTRIBUTIONS

Conceptualization: R.E.J. and A.L.K.; Methodology: R.E.J. and L.A.G.; Validation: R.E.J. and A.L.K.; Formal analysis: R.E.J., N.R.H. and A.L.K.; Investigation: R.E.J., N.R.H., and N.K. Resources: R.E.J., L.A.G., J.P., and A.L.K. Writing – original draft: R.E.J. and A.L.K.; Writing – review & editing: R.E.J., N.R.H., A.L.K, J.P., and L.A.G.; Visualization: R.E.J., N.R.H. Supervision: A.L.K.; Project administration: R.E.J. and A.L.K.; Funding acquisition: R.E.J. and A.L.K.

## DATA AVAILABILITY

scRNA-seq data have been deposited at GEO and are publicly available as of the date of publication. Accession number

All other relevant data can be found within the article and its supplementary information.

